# Single Cell RNA Sequencing Driven Characterization of Pediatric Mixed Phenotype Acute Leukemia

**DOI:** 10.1101/2022.07.07.499210

**Authors:** Hope L. Mumme, Sunil S. Raikar, Swati S. Bhasin, Beena E. Thomas, Deborah DeRyckere, Daniel S. Wechsler, Christopher C. Porter, Sharon M. Castellino, Douglas K. Graham, Manoj K. Bhasin

## Abstract

**Background:** Mixed phenotype acute leukemia (MPAL) is a rare subgroup of leukemia characterized by blast cells that display both myeloid and lymphoid lineage features, making this cancer difficult to diagnose and treat. A deeper characterization of MPAL at the molecular level is essential to better understand similarities/differences to the more common and better-studied leukemias, acute myeloid leukemia (AML) and acute lymphoblastic leukemia (ALL). Therefore, we performed single-cell RNA sequencing (scRNAseq) on MPAL bone marrow (BM) samples in an attempt to develop a more granular map of the MPAL microenvironment landscape.

**Methods:** We analyzed ∼16,000 cells from five pediatric MPAL BM samples collected at diagnosis to generate a single-cell transcriptomic landscape of B/Myeloid (B/My) and T/Myeloid (T/My) MPAL blasts and associated microenvironment cells. Cell clusters were identified using principal component analysis and uniform manifold approximation and projection (UMAP). Unsupervised analysis was performed to determine the overall relationship among B/My MPAL, T/My MPAL, and other acute leukemias – B-ALL, T-ALL, and AML. Supervised differentially expressed gene (DEG) analysis was performed to identify B/My and T/My MPAL blast-specific signatures. MPAL sample transcriptome profiles were compared with normal BM stem and immune cells to identify MPAL-specific dysregulation. Gene set enrichment analysis (GSEA) was performed, and significantly enriched pathways were compared in MPAL subtypes. Comparative analysis was performed on diagnostic samples based on their future minimal residual disease (MRD) and relapse status.

**Results:** B/My MPAL and T/My MPAL blasts displayed distinct subtype-specific blast signatures. UMAP analysis revealed that B/My MPAL samples had greater overlap with B-ALL samples, while T/My MPAL samples clustered separately from other acute leukemia subtypes. Genes overexpressed in both MPAL subtypes’ blasts compared to other leukemias and healthy controls included *PLIN2, CD81*, and *UBE2S*. B/My MPAL blast-specific genes included *IRS2, SMIM3*, and *HBEGF*, whereas T/My MPAL blast-overexpressed genes included *IER5, BOD1L1*, and *HPGD*. Sirtuin signaling, p38 MPAK signaling, and PI3K signaling pathways were upregulated in B/My MPAL blasts while oxidative phosphorylation and Rho family GTPases signaling pathways were upregulated in T/My MPAL blasts. Transcriptomic, pathways, and cell communication level differences were observed in the MPAL samples based on future MRD and clinical outcome status.

**Conclusions:** We have for the first time described the single-cell landscape of pediatric MPAL and demonstrate that B/My and T/My MPAL have unique scRNAseq profiles distinct from each other as well as from ALL and AML.

## Background

Mixed phenotype acute leukemia (MPAL) is a rare subtype of acute leukemia, accounting for 2-3% of all newly diagnosed pediatric leukemia cases, with blasts expressing markers of both the lymphoid and myeloid lineage (1, 2). Antigen expression patterns vary greatly among different MPAL cases, and given the wide phenotypic variability, the diagnostic criteria for MPAL have continued to evolve over the past decades. The European Group for the Immunological Characterization of Leukemias (EGIL) and the World Health Organization (WHO) criteria are the two MPAL classification systems primarily used; however, despite both systems relying on immunophenotypic characterization, there remain significant differences in their definitions (1-4). Given the frequent changes and relative subjectivity in diagnostic criteria, it is extremely difficult to interpret previously published MPAL literature. Reported survival outcomes for MPAL have ranged between 36-80%; however, since patients with an MPAL phenotype were excluded from frontline clinical trials until recently, all available treatment and outcome data for MPAL is retrospective. While most patients with MPAL respond to acute lymphoblastic leukemia (ALL) directed therapy (2, 4), there is no clear consensus on how to treat this heterogeneous disease.

The lack of standardized treatment regimens specifically tailored for MPAL is compounded by fluid diagnostic criteria for classifying MPAL and its subtypes. Current classification systems divide MPAL into two broad categories, B/myeloid (B/My) MPAL and T/myeloid (T/My) MPAL (5). Despite these differences in classification and the wide phenotypic diversity, current treatment approaches have typically considered MPAL to be a single entity with providers primarily choosing between ALL vs. AML regimens, and not considering specific subtypes. Two recent large MPAL genomic studies, one in pediatric patients and one in adults, have shown that B/My and T/My MPAL have distinct genomic signatures (6, 7), supporting the notion that different approaches may be necessary to treat MPAL subtypes. Furthermore, two large retrospective cohorts have now shown that the early response to ALL therapy is critical in terms of overall prognosis (7), with patients having positive measurable/minimal residual disease (MRD) at the end of induction (EOI) having significantly poorer outcomes. Analyzing the biology and understanding the similarities and differences is critical for improving outcomes in these rare high-risk leukemias. Thus, a more in-depth analysis of MPAL biology is essential to determine effective treatments for this unique disease.

ScRNAseq has revolutionized cancer research by revealing cell types, pathways, and cellular interactions that play critical roles in malignant cell progression and response to therapy (8, 9). Identifying changes in cellular and molecular profiles is critical for identifying novel targets for diagnosis, risk assessment, and clinical outcomes. Single-cell profiling can be invaluable for deep characterization, given the wide phenotypic and genomic diversity seen in MPAL. Only one single-cell study of MPAL has been previously reported, using samples from adults (10). Here, we present for the first time scRNAseq profiling of five pediatric MPAL samples collected at diagnosis, along with a comparative analysis with previously generated scRNAseq datasets from pediatric AML and ALL samples as well as young adult healthy bone marrow (BM) samples (11-13). Our analysis revealed similarities between B/My MPAL and B-ALL transcriptomes, while T/My MPAL depicted a unique transcriptome profile distinct from transcriptomes of other leukemias/subtypes. Our results provide an initial framework of the pediatric MPAL single-cell signature and support utilizing scRNAseq analysis for further characterization of the MPAL blast and marrow landscape.

## Methods

### Clinical samples

Primary patient BM samples were obtained from the Aflac Cancer and Blood Disorders Center Biorepository within Children’s Healthcare of Atlanta (CHOA). Patients provided written informed consent that permitted the use of biological material in accordance with a protocol that was approved by the CHOA Institutional Review Board (IRB). The diagnosis of MPAL was made according to the World Health Organization (WHO) 2016 MPAL criteria (5). BM samples were collected at initial presentation as part of routine diagnostic evaluation. The study was performed on five MPAL patient samples (three B/My MPAL and two T/My MPAL immunophenotypes) with known MRD status and clinical outcome (post-therapy remission or relapse). In addition, for comparative analysis, we used single-cell datasets of other pediatric leukemias i.e., AML and T-ALL from previous/ongoing studies in the lab and also, publicly available scRNAseq datasets of young adult healthy BM and pediatric B-ALL and AML samples (13-15).

### Single-cell RNA sequencing and analysis of MPAL samples

Single-cell RNA sequencing (scRNAseq) libraries were prepared from viably revived BM samples using hashtag B antibodies (Biolegend) and Chromium single cell 3’V3 reagent kits (10x genomics). Sequencing was performed using NextSeq 500 high output kits (Illumina) (12). The fastq files were analyzed using Cell Ranger (16) for demultiplexing, alignment to the human genome (hg38), and generation of gene-count matrices for further bioinformatics analysis.

### Single-cell profiling data from other leukemias and healthy bone marrow

For comparative analysis of MPAL with other pediatric leukemias, we used single-cell datasets generated in our lab for other leukemias: AML (n=15), and T-ALL (n=10) (13-15). Data were generated and processed using the uniform approach briefly described in the following paragraph and previously utilized (13-15). Additionally, we also used publicly available datasets, downloaded via the GEO portal (GSE154109), for comparative analysis (17). This dataset contained pediatric B-ALL (n=7), pediatric AML (n=8), and young adult healthy BM (n=4) samples.

### Single-cell profiling data analysis

Raw gene-count matrices from samples were merged to generate a raw expression matrix from all pediatric leukemias and healthy BM. Cells were filtered based on mitochondrial content and feature count (pct. mitochondrial < 60 and feature count < 200). Expression profiles were normalized and scaled using the SCTransform function in Seurat v4 (18). Dimensionality reduction was performed via the UMAP method, and the cells were clustered using the K-nearest neighbor graph-based clustering approach. Leukemic cells, or blasts, were annotated by comparing each leukemia set (AML, B-ALL, T-ALL, MPAL) samples with the healthy control and identified as cells that did not cluster with the healthy control cells. Once the blasts were identified, the non-blast or canonical lymphoid, myeloid, and erythroid lineage cells were annotated based on a combination of automatic annotation using the SingleR package (19), and manual annotation via known marker genes. SingleR is an automatic annotation tool that labels cells based on an external annotated reference, such as the Human Primary Cell Atlas (ERP122984) (20). A supervised analysis was performed to generate MPAL blast transcriptome signatures by comparing gene expression profiles of MPAL subtype blast cells with normal cells using the Wilcoxon rank test. The transcriptome signatures were generated based on fold change and P-value cutoffs (adjusted p-value < 0.05, average log2FC > 0.25, and percent cell expression > 50%). The analysis identified transcriptome signatures for MPAL as well as subtypes i.e., T/My MPAL and B/My MPAL.

### Generation of MPAL-specific gene dysregulation

Transcriptome signatures generated from the previous analysis were systematically compared with normal and stem cell profiles from the human cell atlas (HCA) (21). Blast genes with average expression >0.5 in normal BM or stem cells from the HCA were considered non-specific and filtered out. The analysis resulted in the identification of MPAL blast cell-overexpressed gene sets that were further compared with pediatric ALL and AML leukemia blast cells to identify MPAL specific gene sets with high potential to be MPAL biomarker candidates (**Fig. S1**).

A similar differentially expressed gene (DEG) and biomarker analysis was performed on the future relapse and remission blast cell profiles to identify genes that are specifically overexpressed in MPAL remission- or relapse-associated blasts.

### Pathway enrichment analysis

MPAL blast-specific pathways were identified by performing pathway analysis using the Ingenuity Pathway Analysis software (IPA) by QIAGEN Inc. (22). The B/My MPAL blast-specific gene set was generated by identifying significantly DEGs between B/My MPAL blasts, other leukemia blasts, as well as immune cells (p-value <0.05 and log2FC >0.25). The same process was performed to identify a T/My MPAL blast-specific gene set. A detailed description of IPA is available on the QIAGEN website (23). IPA calculates a p-value for each pathway according to the fit of the user’s data to the IPA database using a one-tailed Fisher exact test. Pathways with p-values <0.05 were considered significantly affected.

Additionally, we also performed pathway and systems biology analysis on MRD-associated genes using the MetaCore platform (Clarivate Inc.). The MRD positivity-associated signature was identified by comparing blast cells from MRD positive and negative samples by linear statistical analysis using the Wilcoxon rank test (p-value<0.05, absolute log2FC>0.25). The list of DEGs was submitted to the platform for analysis. The knowledge base of this platform consists of functions, pathways, and network models derived by systematically exploring peer-reviewed scientific literature and public databases. It calculates statistical significance based on the hypergeometric distribution where the p-value represents how likely the observed association between a specific pathway/function/interactive network and the dataset would be if it were only due to random chance, by also considering the total number of functions, pathways, and interactive network eligible genes in the dataset and the Reference Set of genes (those which potentially could be significant in the dataset). Focus molecules were identified from the integrated networks based on the degree of connectivity (number of interactions for each gene). Focus hubs with higher degrees of connectivity are considered critical for the maintenance of the networks, suggesting that therapeutic targeting of these focus hubs may elicit the strongest impact. Pathways and networks with a p-value <0.05 were considered statistically significant.

### Gene-set enrichment analysis

In addition to individual gene analysis, gene set enrichment analysis (GSEA) was implemented to determine whether an *a priori*-defined set of genes showed statistically significant, concordant differences between different group comparisons (24). GSEA can be more powerful than single-gene methods for studying the effects of factors such as MRD in which many genes each make subtle contributions. GSEA was performed using the escape R package (25). Gene sets with a p-value <0.05 were considered significantly altered. GSEA was performed on the basis of 3,000 canonical pathways obtained from the MSig database via the msigdbr package (v7.4.1). Once the significantly upregulated pathways were identified for each leukemia type, the comparative analysis of pathways resulted in T/My and B/My MPAL-specific pathways from blast cells as well as major immune cell subtypes including “B, Pro-B, and plasma cells”, “T and NK cells”, “progenitor cells”, and “monocytes and macrophages”.

### Cellular communication analysis

Cellular communication analysis was performed using CellChat version 1.0.0 (26). CellChat uses ligand-receptor expression to predict intercellular communication among specific signaling pathways. CellChat first calculates the communication probability of each signaling pathway in the CellChatDB ligand-receptor database between cell clusters or cell types within a group (e.g., diagnostic sample from a patient who relapsed; Dx-Rel). Differences in cellular communication between groups can be analyzed by calculating the information flow, the sum of communication probability for each cell cluster predicted interaction, for each signaling pathway. Signaling network differences between groups can also be analyzed by performing manifold learning and classification based on functional similarity, or the similarity in sender/receiver cell types between two pathways. CellChat also allows analysis of specific cell type interactions within each group for each signaling pathway, and this can be visualized on a circular chord diagram.

### Survival analysis

Estimated survival probabilities for the T-Myeloid MPAL and B-Myeloid MPAL Dx-Rel and Dx-Rem (diagnostic samples from patients that underwent remission) biomarker sets were calculated using the survMisc (27) and survival (28) R packages. Survival analysis was performed on the TARGET-ALL-P3 dataset (ambiguous leukemias) after extracting the T-Myeloid and B-Myeloid MPAL samples (29). The T-Myeloid MPAL Dx-Rel and Dx-Rem gene sets were used to calculate gene set enrichment values for each sample with a T-Myeloid MPAL diagnosis. Then, the samples were split into high and low expression groups using the cutP method (27) based on the samples’ gene set enrichment values for each gene set. The survival was calculated, and Kaplan-Meier survival curves and hazard ratio statistics were produced. The same analysis was also performed for the B-Myeloid MPAL gene sets.

## Results

### Characterizing the single-cell landscape of MPAL BM samples

To characterize the tumor microenvironment (TME) of MPAL, we performed scRNAseq on BM samples of three B/My MPAL and two T/My MPAL patients, all of whom were treated with an ALL induction regimen (**Table 1)**. Of three B/My MPAL samples, two were MRD+ (M3, M5) and one was MRD- (M1), while both T/My MPAL samples (M4, M6) were MRD+ at EOI (**Table 1**). One B/My MPAL patient relapsed post-therapy and two achieved remissions while one T/My MPAL patient relapsed and one achieved remission post-therapy (**Table 1**).

**Table 1:** MPAL patient clinical information. Patient characteristics and clinical information with sample ID, leukemia subtype diagnosis, initial white blood cell (WBC) count, clinical peripheral blood (PB) blast percentage, clinical BM aspirate (BMA) blast percentage, World Health Organization (WHO) MPAL immunophenotyping markers, minimal residual disease (MRD) status (< 0.01% classified as MRD negative, > 0.01% classified as MRD positive) at end of induction (EOI), and clinical status (relapse vs remission).

| Sample ID | Diagnosis | Initial WBC (x103/uL) | PB Blast (%) | BMA Blast (%) | WHO MPAL markers | EOI MRD Status | Patient Relapsed |
| --- | --- | --- | --- | --- | --- | --- | --- |
| M1 | B/My MPAL | 134.6 | 95.7 | 97.7 | B-lineage: CD19, CD22; Myeloid: CD11c, CD64 | Negative | No |
| M3 | B/My MPAL | 49.97 | 31.8 | 82.0 | B-lineage: CD10, CD19, CD22; Myeloid: CD11c, MPO | Positive | Yes |
| M4 | T/My MPAL | 33.03 | 73.5 | 81.4 | T-lineage: CD3; Myeloid: CD11c, CD64 | Positive | Yes |
| M5 | B/My MPAL | 12.83 | 48.0 | 94.4 | B-lineage: CD10, CD19, CD22; Myeloid: CD14, CD64, MPO | Positive | No |
| M6 | T/My MPAL | 7.1 | 5.0 | 40.2 | T-lineage: CD3; Myeloid: CD11c, CD64, MPO | Positive | No |

Comprehensive single-cell transcriptome profiling was performed using the 10x genomics platform after reviving viably frozen patient BM samples collected at the time of disease diagnosis (Dx). To identify blast cells, the data from healthy young adult BM samples were included in the analysis (17). In total, we analyzed 22,348 cells (5,443 from two T/My MPAL, 10,961 from the three B/My MPAL, and 5,944 from healthy BM samples). After undergoing quality control and normalization, unsupervised analysis identified 20 transcriptionally distinct clusters of cells (**Fig. 1A**). Most cellular clusters were labeled based on the expression of canonical cell lineage-associated markers (**Fig. S2**); these cell types include erythroid, monocyte/macrophage, T-cells, and cytotoxic T lymphocytes (CTL), B-cells, Pro-B, NK, and general myeloid progenitors (GMP). The clusters (M1, M3, M4, M5, and M6 blasts) lacking canonical immune cell markers and predominantly made up of cells from individual patients were considered putative blast cell clusters. The putative MPAL leukemic blast clusters depicted segregated clustering from the healthy control cell clusters (**Fig. 1B**, lassoed regions). B/My MPAL blasts formed patient-specific segregated clusters (M1, M3, M5 blasts) indicative of inter-patient heterogeneity (**Fig. 1B**) while the T/My MPAL blasts (M4, M6 clusters) showed inter-patient similarity with overlapping clusters (**Fig. 1B**). On the other hand, immune cells from both MPAL subtypes clustered together with no inter-patient heterogeneity (**Fig. 1B**). The analysis of MPAL and healthy BM samples based on UMAP depicted that MPAL BM microenvironment cells also have subtle differences as compared to corresponding healthy cells. To identify the top differences between MPAL microenvironment cells as compared to healthy immune cells, we performed a supervised differential expression analysis based on the Wilcoxon rank test (**Fig. 1C**). The MPAL B cells overexpressed *IGHG1, JCHAIN*, and *IGHG3* as compared to healthy B-cells, indicating enrichment of plasma cell lineage in the MPAL samples. Similar supervised analysis of cytotoxic T cells depicted significant overexpression of *NFKBIA*, and *COTL1*, indicating enrichment of T-cell activation regulators (30, 31). Lists of the top 10 upregulated genes in MPAL vs. healthy samples in non-blast clusters are available in **Table S1**. There were no significant differences in cell type abundances between B/My MPAL and T/My MPAL; however, there were significant differences in healthy and MPAL cell type abundances in B cells, T cells, and Pro-B cells (**Fig. 1D**). This reduction in immune cells in the MPAL bone marrow microenvironment is likely due to marrow infiltration by the malignant blast cells.

**Figure 1:**
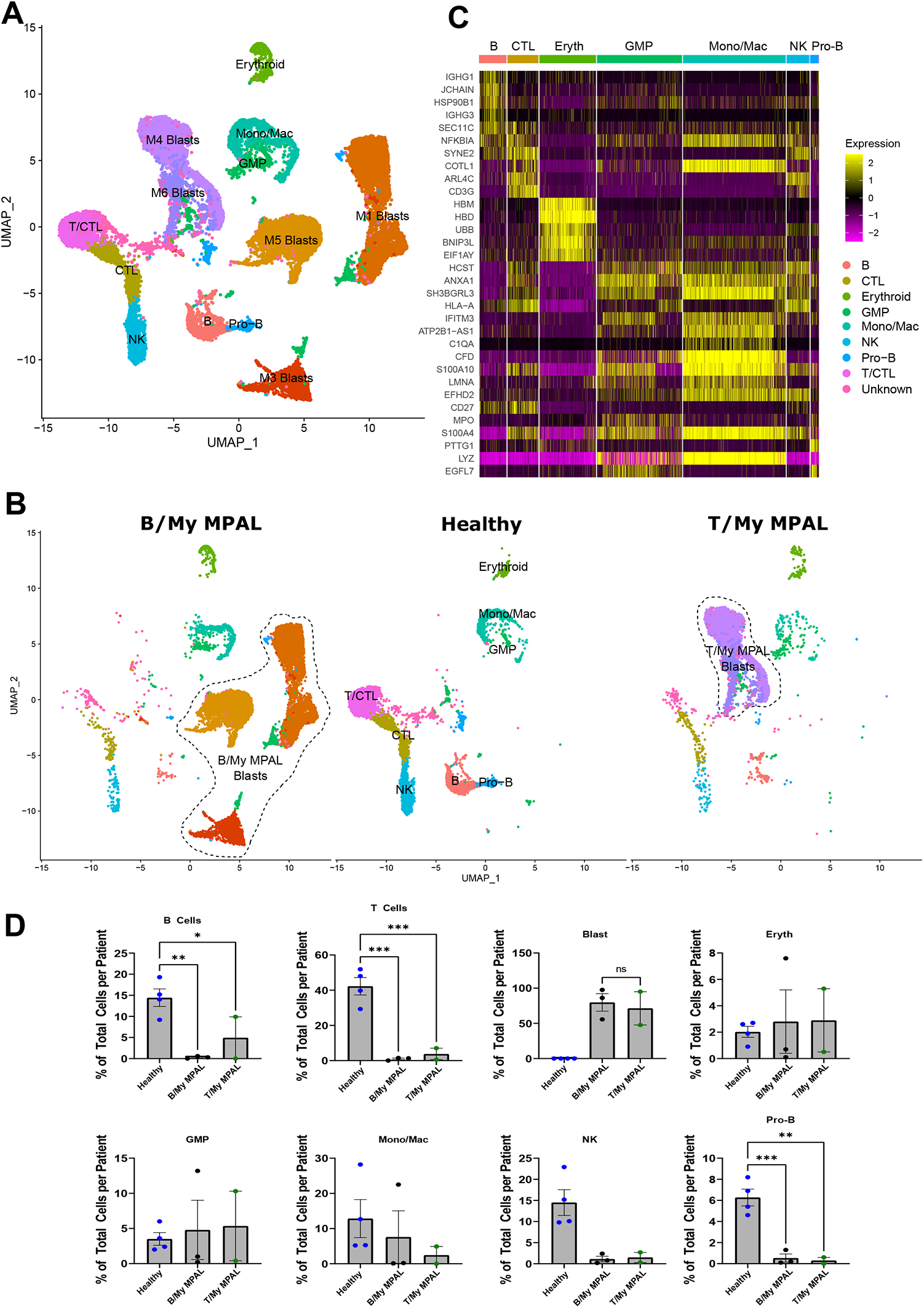
Single-cell landscape of MPAL and healthy control bone marrow samples. **A)** Uniform manifold approximation and projection (UMAP) embedding of the MPAL and healthy control samples consisting of 22,348 cells. The cells are colored by distinct cell type clusters and labeled manually based on overexpression of cell type-specific markers (T-lymphocytes (T), cytotoxic T cells (CTL), B-lymphocytes (B), progenitor B cells (Pro-B), natural killer cells (NK), monocytes/macrophages (Mono/Mac), erythroid, granulocyte-monocyte progenitor (GMP), and patient-specific clusters). **B)** Split UMAP based on the clinical groups (B/My-MPAL, Healthy, T/My-MPAL). T/My-MPAL and B/My-MPAL clinical group-specific clusters are marked with a lasso. **C)** Heatmap showing the top highly expressed (yellow) genes in each of the MPAL immune cell types in comparison to corresponding healthy cell types. **D)** Bar plots showing the percentage of various immune cells across three clinical groups (Healthy, T/My-MPAL, and B/My-MPAL). The healthy control samples have a significantly higher percentage of adaptive and innate immune cells as compared to both MPAL sub-types. Results are expressed using bar graphs representing the mean and SEM values in the groups (*p-value < 0.05, ** p-value < 0.01 by group-wise comparisons using ordinary one-way ANOVA).

### B/My MPAL scRNAseq profile has significant overlap with B-ALL, whereas T/My MPAL has a unique profile

To assess the similarities and differences between MPAL and other leukemias, we performed comparative analyses among MPAL, AML, B-ALL, T-ALL, and healthy BM single-cell profiles. Single-cell transcriptome data for other leukemias were obtained from the Pediatric Cancers Single-Cell Atlas (15) initiative of our lab. After uniform pre-processing, filtering, and normalization, the average gene expression of each sample was compared using UMAP and principal component analysis (PCA) (**Fig. 2**). UMAP analysis showed the B/My MPAL blasts are clustered with B-ALL, indicating transcriptional similarity between B/My MPAL and B-ALL (**Fig. 2A, B**). The T/My MPAL blasts, however, clustered distinctly from T-ALL and other leukemias, indicating a unique T/My MPAL transcriptional landscape (**Fig. 2A, B**). Interestingly, a minor fraction of T-ALL cells (∼2% of cells) was observed in the T/My MPAL cluster, suggesting the existence of rare T/My like blasts in the T-ALL. The 3D PCA analysis also depicted that T/My MPAL samples are segregated from other leukemias as well as healthy BM controls. On the other hand, B/My MPAL samples were shown to have a profile overlapping with B-ALL (**Fig. 2C**).

**Figure 2:**
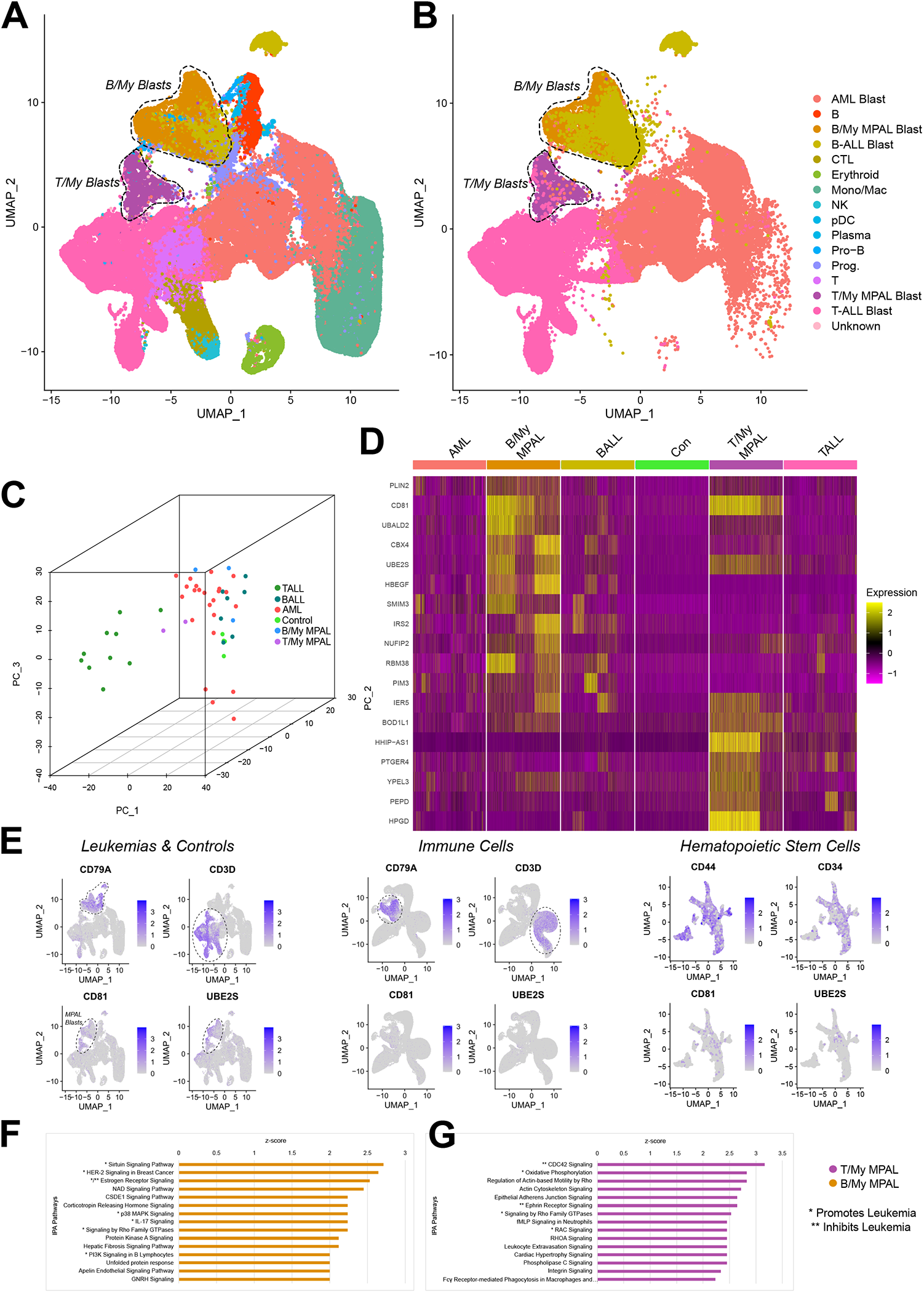
Comparative analysis of MPAL blast cells with other leukemias and healthy bone marrow. **A)** Uniform manifold approximation and projection (UMAP) embedding of the AML, B-ALL, T-ALL, and healthy bone marrow samples consisting of 122,376 cells. The cells are colored by distinct cell type clusters and labeled manually based on overexpression of cell type-specific markers (T-lymphocytes (T), cytotoxic T cells (CTL), B-lymphocytes (B), progenitor B cells (Pro-B), natural killer cells (NK), monocytes/macrophages (Mono/Mac), erythroid, granulocyte-monocyte progenitor (GMP), AML blasts, T-ALL blasts, B-ALL blasts, T/My MPAL blasts, and B/My MPAL blasts). **B)** UMAP embedding of the AML, B-ALL, T-ALL, T/My MPAL, and B/My MPAL blast cells, showing overlap of B/My MPAL with B-ALL and unique profile for T/My MPAL. **C)** Principal component analysis (PCA)-based comparison of average transcriptome profile of MPAL, AML, B-ALL, T-ALL, and healthy BM samples. **D)** Heatmap showing significantly overexpressed genes among MPAL blast cell clusters compared to blast cells from other leukemias (AML, B-ALL, T-ALL) and healthy BM. Relative gene expression is shown in pseudo color, where pink represents low expression, and yellow represents high expression. **E**) Feature plots showing the expression of MPAL blast-specific genes in pediatric leukemias and healthy BM control, HCA (21) immune cell, and HSC datasets. MPAL marker genes (*CD81, UBE2S*) have a significant expression in the blast cell clusters and minimal expression in immune cell clusters which showed overexpression of immune cell marker genes (*CD79A, CD3D*), and in stem cell clusters which showed overexpression of HSC marker genes (*CD44, CD34*). **F, G)** Pathways enrichment analysis of genes significantly overexpressed (p-value < 0.05, average log2FC > 0.5) in MPAL blast cells. B/My and T/My MPAL significantly upregulated pathways based on Z score and P-value are shown in orange and purple colors, respectively. Pathways with evidence for promoting (*) or inhibiting (**) leukemia are marked with one or two asterisks, respectively.

To identify genes with significant overexpression in MPAL as compared to other leukemias and healthy controls, DEG analysis was performed based on the Wilcoxon rank test (FC > 1.2 and adjusted P-value < 0.05). Genes that are overexpressed in both MPAL subtypes’ blasts compared to other leukemias and healthy controls include *PLIN2, CD81*, and *UBE2S* (**Fig. 2D**). B/My MPAL blast genes include *IRS2, SMIM3*, and *HBEGF*. T/My MPAL blast genes include *IER5, BOD1L1*, and *HPGD*. To ensure the MPAL blast markers are tumor-specific, a few additional filtering steps based on expression in the BM immune cells and hematopoietic stem cells (HSCs) were implemented. The BM single-cell data of 391,505 immune cells and HSCs were obtained from the Human Cell Atlas (HCA) Initiative (21). MPAL blast-specific genes had minimal expression in the HCA immune cells and the HCA HSCs (**Fig. 2E**), whereas canonical immune cell markers such as B cell gene *CD79A* and T cell gene *CD3D* had high expression in the HCA immune cells and well-known HSC marker genes such as *CD44* and *CD34* had high expression in the HSCs. The complete lists of MPAL blast markers with average log2 fold change and p-values from each step of the filtering process (**Fig. S1**) are listed in **Table S2**. The biomarker genes (**Fig. 2D**) had minimal or no expression in normal immune and stem cells (**Fig. S3**), making these genes ideal candidates to be explored for developing targeted therapies to improve MPAL outcomes.

To further understand the biological pathways level of dysregulation in MPAL blast genes, pathways enrichment analysis was performed on the top significantly expressed genes (p-value <0.05, average log2FC >0.5). Sirtuin signaling, p38 MPAK signaling, and PI3K signaling pathways were upregulated in B/My MPAL blasts and have been found to promote leukemia cell survival in other types of leukemia (32-34) (**Fig. 2F**). In T/My MPAL blasts, oxidative phosphorylation, and Rho family GTPases signaling pathways were upregulated (**Fig. 2G**) and have been shown to promote leukemia cell survival (35). Common pathways that were significantly impacted in both B/My and T/My MPAL included ILK signaling (36), ERK/MAPK signaling (37), and PI3K/AKT signaling (38) (**Fig. S4**). The complete lists of significantly affected pathways with p-values and Z-scores are shown in **Table S3**.

### MPAL microenvironment cells display specific dysregulations at the pathway and cellular communication network levels

Cellular communications between cell types in the MPAL subtypes were compared against cell communication in other leukemias and controls (**Fig. S5**). For each signaling pathway found to be enriched in the MPAL subtypes, the analysis identified cell types and their levels (i.e., heatmap color intensity) of involvement in the incoming (receptor) or outgoing (ligand) signaling (**Fig. 3A**). Of the signaling pathways that were estimated to be present in B/My MPAL samples, the neuronal growth receptor (NEGR) signaling pathway was significantly enriched in B/My MPAL **(Fig. S5)** but not in other leukemias or control groups. The NEGR pathway had incoming and outgoing signaling exclusively present in Pro-B cells (**Fig. 3A**). Similarly, T/My MPAL blasts demonstrated the strongest outgoing signal in the TGF beta pathway, which was specifically enriched in T/My MPAL samples, with Pro-B cells being the receiving cell type. Overall, the CD99, MHC-II, MIF, GALECTIN, CLEC, and MHC-I pathways had the highest information flow among the enriched pathways of B/My MPAL. Among the T/My MPAL pathways, the MIF, CLEC, and MHC-I pathways had the highest information flow (**Fig. 3A**).

**Figure 3:**
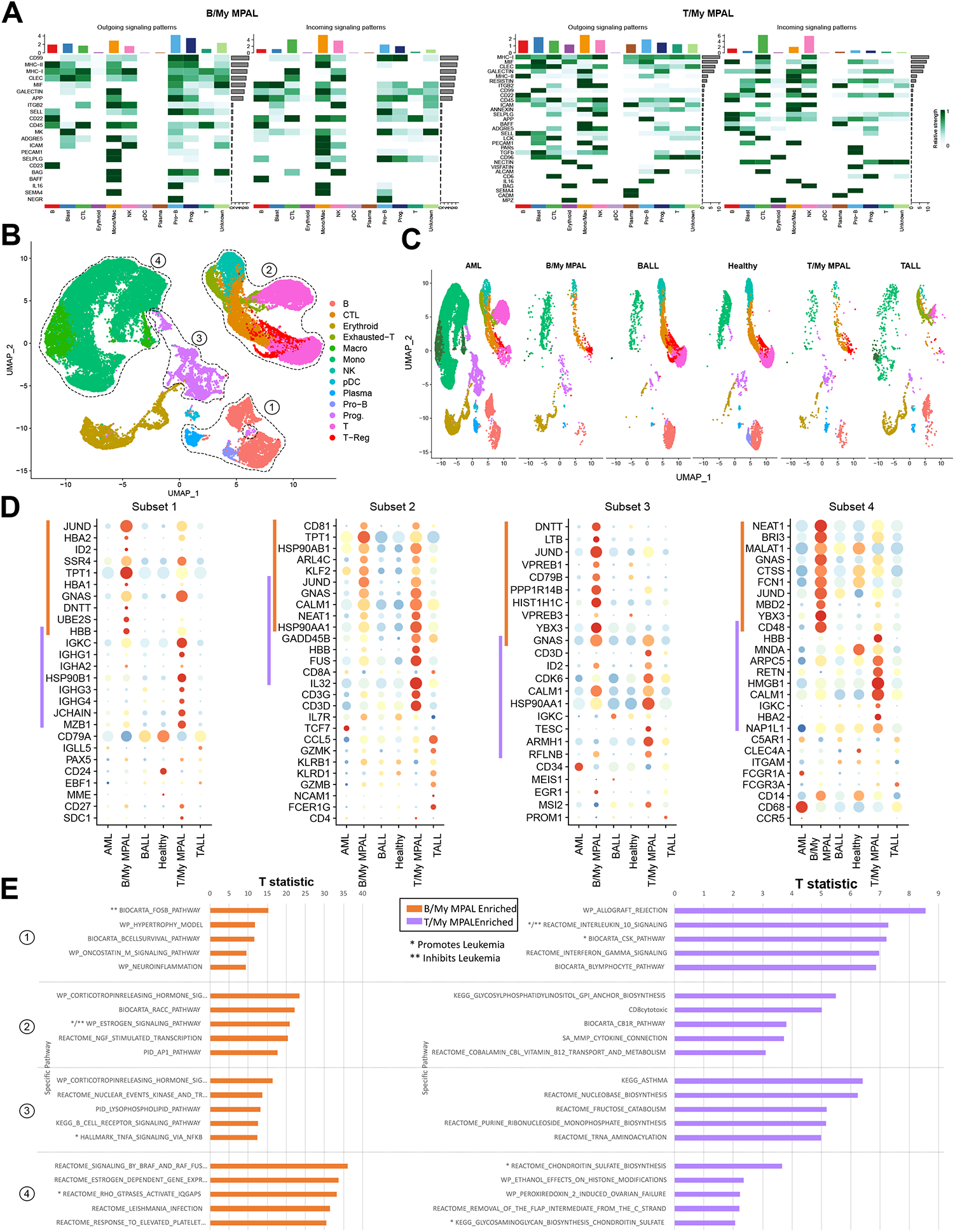
Cellular communications and pathways analysis of MPAL microenvironment cell lineages. **A)** Cellular communication analysis showing top incoming and outgoing cellular pathway enrichments in various B/My and T/My MPAL cell types. The relative strengths of incoming (receptor) or outgoing (ligand) signaling pathways are represented in different shades of green color. The overall cellular communication activity of different cell types is shown with bar graphs attached to the heatmap. **B)** UMAP embedding of the microenvironment cell types of MPAL, AML, B-ALL, T-ALL, and healthy BM. The individual cell types were colored by distinct cell type clusters and labeled manually based on overexpression of cell type-specific markers (T-lymphocytes (T), cytotoxic T cells (CTL), B-lymphocytes (B), progenitor B cells (Pro-B), natural killer cells (NK), monocytes/macrophages (Mono/Mac), erythroid, and granulocyte-monocyte progenitor (GMP)). The four major lineages of cell types are marked with the lasso: subset 1; B-lymphoid (B-, pro-B-, and plasma-cells), subset 2; T/NK-lymphoid (exhausted-T-, T-, T-reg, and NK-cells), subset 3; progenitor cells, and subset 4; myeloid lineage (monocytes and macrophages). **C)** Split UMAP based on clinical groups showing relative enrichment of cell types. **D)** Dot plots of top MPAL overexpressed genes across four cell lineages. The canonical cell type markers for the different cell lineages are also included in the plots for cellular annotations. **E)** Gene set pathway enrichment analysis across four cell lineages analysis to identify the top significantly dysregulated pathways in B/My and T/My MPAL. B/My and T/My MPAL significantly enriched pathways T-statistics are shown in orange and purple colors, respectively. Pathways with evidence for promoting (*) or inhibiting (**) leukemia are marked with one or two asterisks, respectively.

To further characterize transcriptome and pathway level differences among microenvironment cell types of different leukemias including MPAL, and healthy BM, we performed a focused analysis of microenvironment cell types only. **Fig. 3B** shows the annotated UMAP of immune and other microenvironment cells, with four major lineages: subset 1: B-lymphoid (B-, pro-B-, and plasma-cells), subset 2: T/NK-lymphoid (exhausted-T-, T-, T-reg, and NK-cells), subset 3: progenitor cells, and subset 4: myeloid (monocytes and macrophages). Comparative analysis of different leukemias (AML, B-ALL, T-ALL, B/My MPAL, and T/My MPAL) and healthy BM revealed that most cell types are detected across all groups with subtle differences in enrichment (**Fig. 3C**). DEG analysis comparing expression profiles among MPAL subtypes vs. other leukemias, and healthy BM defined key B/My (shown by orange bars) and T/My (shown by purple bars) MPAL-specific markers for different cell types **(Fig. 3D)**. The B-lymphoid lineage (subset 1) of B/My MPAL shows specific overexpression of cell proliferation and differentiation genes including *JUND, DNTT*, and *ID2*, whereas T/My MPAL shows overexpression of immunoglobin genes including *IGHG1, IGHA2*, and *JChain* **(Fig. 3D)**. The T/NK-lymphoid lineage (subset 2) shows a significantly similar transcriptome profile between T/My and B/My subtypes with overexpression of genes like *CALM1* and *NEAT1* **(Fig. 3D)**. *NEAT1* plays a key regulatory role in T cell function and could be an important target for enhancing the efficiency of immunotherapy (39, 40). The progenitor lineage (subset 3) of B/My and T/My MPAL shows specific overexpression of *YBX3* and *CDK6*, respectively, which are key regulators of gene expression and cell cycle regulation (41, 42) (**Fig. 3D**). The canonical cell type markers for the different lineages are also included in the four plots in **Fig. 3D**.

To further characterize MPAL microenvironment-associated transcriptome differences, we performed pathway level analysis using the gene set enrichment approach (43, 44). The pathways for B/My MPAL and T/My MPAL only included those that were specific to the subtype and not upregulated in any other leukemia (**Fig. 3E**). Among the top unique pathways enriched in B/My MPAL, T-lymphoid lineage cells were those involved in corticotropin-releasing hormone (CRH) signaling, nerve growth factor (NGF) signaling, and activator protein-1 (AP1) (**Fig. 3E**). The B/My MPAL B-lymphoid lineage demonstrated significant enrichment of FOSB pathways, which has been associated with poor outcomes in AML (45). The TNFA signaling pathway via NF-κB was also found to be specifically enriched in a progenitor lineage that is known to promote leukemia cell survival in AML (**Fig. 3E**) (46). The B-lymphoid lineage of T/My MPAL showed significant enrichment of the IL-10 pathway that has been extensively explored as a prognostic marker for leukemias (47). Also, in T/My MPAL B cells, the upregulated CSK pathway is associated with the promotion of B-cell activation, while the IFN-γ pathway is associated with inhibition of B-cell activation (**Fig. 3E**) (48).

### Comparative analysis of diagnostic B/My MPAL samples based on future EOI MRD status reveals distinct blast cells associated with expression profiles

To explore the association between MRD status of the blast cells and transcriptome profile at baseline, we performed a focused analysis of B/My MPAL MRD positive vs. negative blast cells. There were two MRD positive and one MRD negative sample among the B/My MPAL samples. Clustering and UMAP embeddings based on MRD status revealed distinct blast cell transcriptome profiles (**Fig. 4A**). Blast cells from each sample depicted heterogenous transcriptome profiles evident from patient-specific clusters whereas non-blast cells (B, CTL, Erythroid, GMP/Prog, Mono/Mac, NK, Pro-B, T/CTL) formed mostly overlapping clusters (**Fig. 4A**). MRD positivity associated genes were identified through differential expression analysis by comparing the profile of blast cells from MRD positive and negative samples based on the Wilcoxon rank test. The analysis identified 609 genes that were significantly differentially expressed, 95 upregulated and 514 downregulated in MRD-positive samples (**Fig. 4B)**. To identify the key pathways and regulators affected by the MRD positive blast-associated genes, we performed a pathways and systems biology-oriented analysis. Interestingly, antigen presentation and inflammatory response-related pathways emerged as the top pathways that were significantly affected in the MRD positive group (**Fig. 4C**). The induction of antigen presentation machinery by MHC I was significantly more activated in the MRD negative compared to MRD+ samples, indicative of the critical role of the immune response in eradicating the disease (**Fig. 4C**). Further network analysis identified significant activation of regulatory immune networks related to IRF1 (**Fig. 4D**), MHC Class II, LAPTM5, and C-Fos (**Fig. 4E**), suggesting their role in MRD positivity.

**Figure 4:**
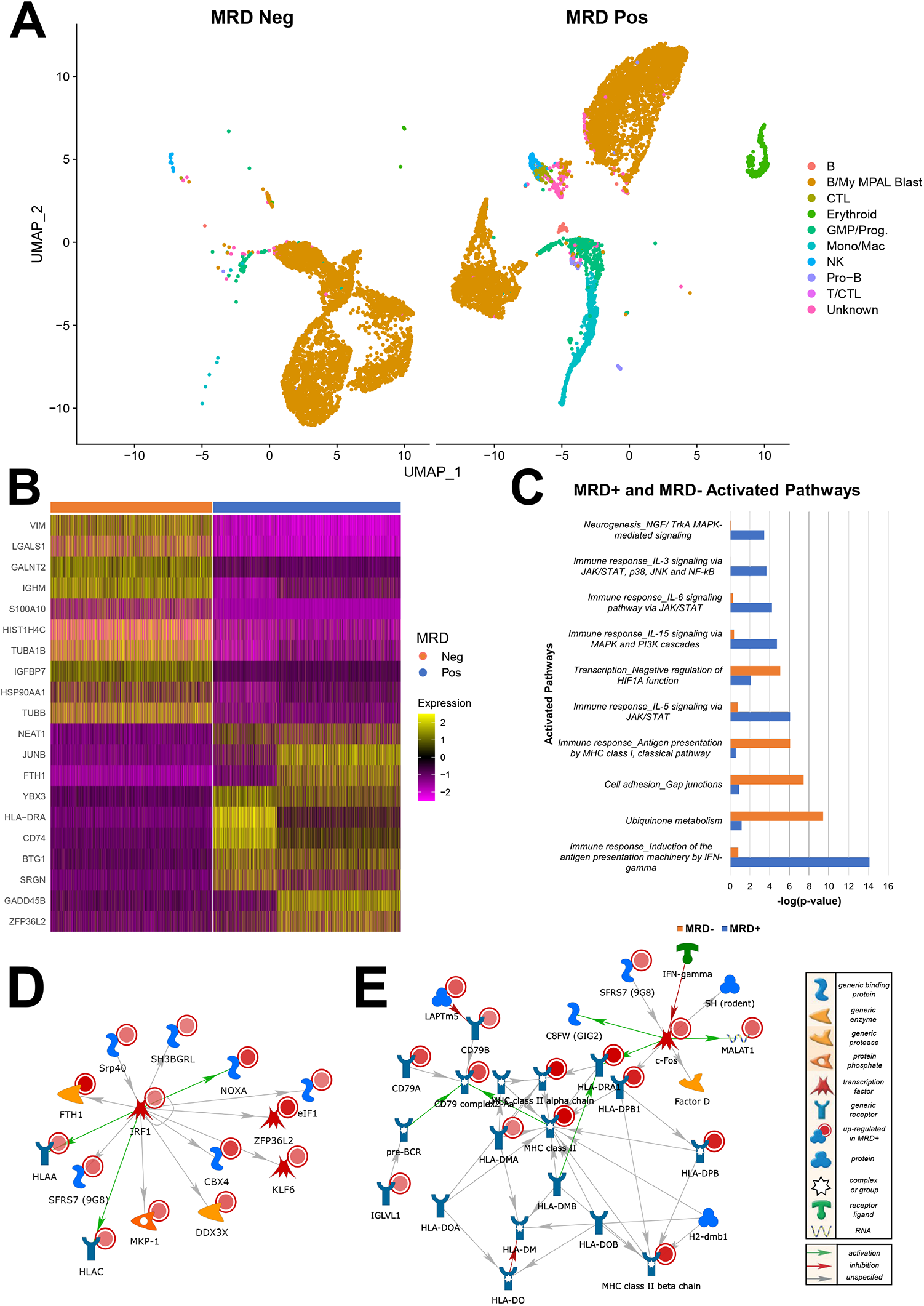
Transcriptomics differences in MPAL blast cells based on MRD status. **A)** Split UMAP of blast and microenvironment cell types based on future MRD positive and negative MPAL samples. Blasts and different microenvironment cells were annotated based on blasts and canonical cell markers identified in **Fig. 1. B)** Heatmap of top genes that are significantly differentially expressive in MRD positive vs negative samples. In the heatmaps, columns and rows represent the cell types and genes, respectively. The yellow color represents high average gene expression while pink represents low gene expression. **C)** Pathways that are significantly affected based on DEGs, i.e., significantly upregulated (Blue) and downregulated (Orange) in MRD positive vs negative samples. Pathways achieved a p-value <.05 based on the hypergeometric distribution where the p-value represents the probability of mapping between a gene set and pathways due to random chance. **D)** IRF1 regulated interactive network that is significantly activated in MRD positive samples. Each node represents a gene and edges interactions among them. **E**) Top network that is significantly activated in MRD positive samples with MHCII and C-Fos as master regulator genes. The pathways and network analysis were performed using the MetaCore tool (Clarivate Inc). The legend on the right shows the description of each of the symbols in parts **D** and **E**, with red circles indicating that the particular molecule was found to be upregulated in MRD positive samples.

### Transcriptomic differences in MPAL blast cells based on future relapse or remission status

MPAL subtype diagnostic samples were split into future relapse (Dx-Rel) and remission (Dx-Rem) groups based on the clinical outcome; there were two B/My MPAL Dx-Rem samples and one Dx-Rel sample, and one of each group for T/My MPAL. The comparative analysis of Dx-Rel and Dx-Rem groups based on UMAP embedding showed differences in the cellular enrichment as well as transcriptome profile (**Fig. 5A**). To identify genes showing dysregulation between blast cells of relapse and remission groups, we performed DEG analysis on individual MPAL subtypes and outcomes (i.e., Dx-Rel, Dx-Rem) (**Fig. 5B**). To identify markers that are specifically dysregulated in relapse/remission blast cells, we also performed a comparative analysis with HCA (21) healthy bone marrow cells and stem cells to filter out genes that are ubiquitously expressed. For B/My MPAL, Dx-Rel markers included *ECM1, SPATS2L*, and *HLA-DQA2*, and Dx-Rem markers included *ATF3, MKNK2*, and *RBM38*. For T/My MPAL, Dx-Rel markers included *HPGD, KRT1*, and *HHIP-AS1*, and Dx-Rem markers included *CDKN2A* and *GLS*. These genes associated with outcomes (**Fig. 5B**) represent potential prognostic blast markers for MPAL subtypes and can be explored in the future as potential therapeutic targets. Further, to understand pathway level differences among Dx-Rel and Dx-Rem groups for both MPAL subtypes, we performed pathways analysis (**Fig. 5C**). For B/My MPAL, Dx-Rel enriched pathways included T cell receptor signaling, the Th1 pathway, and NFAT involvement in the immune response, and Dx-Rem enriched pathways included PD-1 and PD-L1 immune checkpoints, BAG2 signaling, and p38 MAPK signaling. For T/My MPAL, Dx-Rel enriched pathways included CDC42 signaling, Rho regulation of actin-based mobility, and TNFR1 signaling, and Dx-Rem enriched pathways include EIF2 signaling, protein kinase A signaling, and ERK5 signaling.

**Figure 5:**
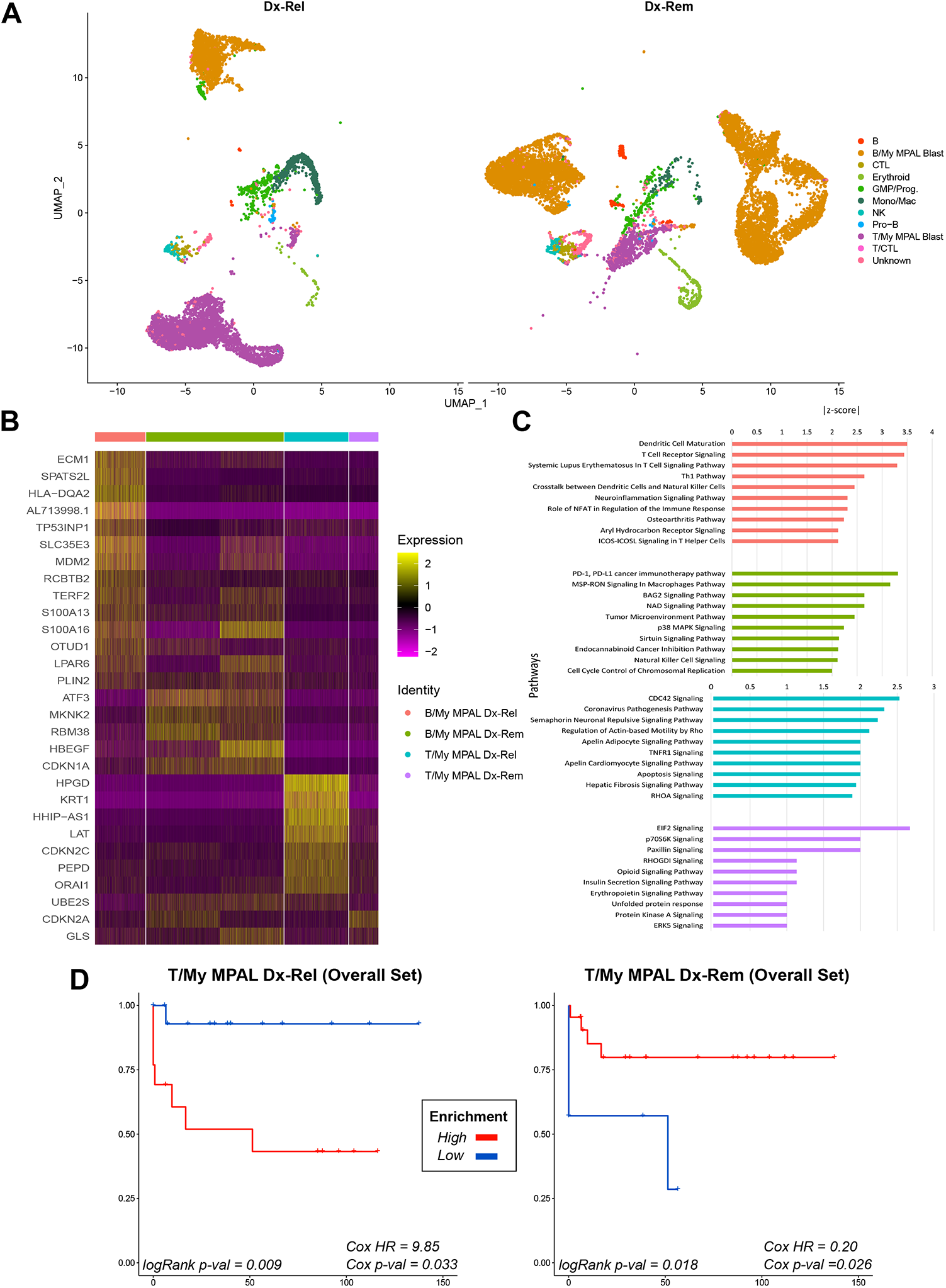
Transcriptomics differences in MPAL blast cells based on outcomes (i.e., future relapse or remission). **A)** Split UMAP of blast and microenvironment cell types from future relapse (Dx-Rel) and future remission (Dx-Rem) samples. Blast and different microenvironment cells were annotated based on blast and canonical cell markers identified in **Fig. 1. B)** Dx-Rel and Dx-Rem blast markers for both MPAL subtypes, identified using differential gene expression analysis and filtering with HCA (21) immune cells and HSCs. **C)** Dx-Rel and Dx-Rem blast-activated pathways for both MPAL subtypes, identified using IPA-based pathways enrichment analysis. The pathways with multiple tests corrected p-value <.05 and Z score >1 were considered significantly activated. **D)** Survival association of T/My MPAL remission and relapsed associated gene signatures (log2FC > 1.25 and adjusted p-value < 0.01). Kaplan Meier plots show high expression of T/My MPAL relapse-associated genes were associated with poorer OS in the TARGET acute leukemias of ambiguous lineage (ALAL) dataset. Similar analysis of T/My genes with greater expression in the remission enriched blast cells showed a significant association with better OS.

To explore the survival association of the top overexpressed genes (log2FC >1.25 and adjusted p-value <0.01) in Dx-Rel and Dx-Rem MPAL samples, we used external bulk RNA-seq data of acute leukemias of ambiguous lineage (ALAL) from the TARGET initiative. The TARGET Phase III dataset includes expression profiles for 115 pediatric leukemia ALAL patients generated using the bulk RNA sequencing approach. The B/My MPAL (n = 25) and T/M MPAL (n = 29) cases were extracted from the TARGET dataset for this analysis. The Cox hazard ratio (HR), Cox p-value, and logrank p-value were determined using the cutP method (27) for each gene set (n = 25 genes for B/My MPAL Dx-Rel, n = 14 for B/My MPAL Dx-Rem, n = 12 for T/My MPAL Dx-Rel, and n = 15 for T/My MPAL Dx-Rem, **Table S4**). For B/My MPAL, the Dx-Rel overall gene set included S100A8, HLA-DRB5, and FCN1, and the Dx-Rem set included IGHM, NR4A1, and IER2. For T/My MPAL, the Dx-Rel overall gene set included HES4, GSTP1, and KRT1, and the Dx-Rem gene set included ACTG1, RACK1, and NEAT1. Of the four sets of genes, only the sets for T/My MPAL Dx-Rel and Dx-Rem had significant survival associations (Cox p-value <0.05) (**Fig. 5D**). The T/My MPAL Dx-Rel genes had a Cox HR of 9.85 (Cox p-value = 0.033, logRank p-value = 0.009), the Dx-Rem genes had a Cox HR of 0.20 (Cox p-value = 0.026, logRank p-value = 0.018). The B/My MPAL outcome-associated genes demonstrated no significant association with outcome (**Fig. S7**).

### Comparison of MPAL microenvironment cells and communication networks based on clinical outcomes

To characterize the MPAL microenvironment in future remission (Dx-Rem) and relapse (Dx-Rel) samples, we performed a cellular communication analysis based on the expression of ligand and receptor pairs among various cell types. The overall cellular communication of MPAL was estimated and the number of signaling interactions was quantified for each cell type in the Dx-Rel and Dx-Rem groups (**Fig. 6A**). Due to the limited number of non-blast cells for each cell type present in the diagnostic samples, the MPAL subtypes were combined for this stage of the analysis. The future remission subset (Dx-Rem - Blue) had a higher number of interactions involving the GMP, Mono/Mac, NK, Pro-B, and T cell types as compared to the future relapse subset (Dx-Rel - Red). The Dx-Rel subset had a higher number of interactions between the blasts and B cells, as well as GMP and NK to blast cells. The higher number of B cell interactions in the future relapse samples might be due to the difference in the number of B cells in the Dx-Rel (n = 10) and Dx-Rem (n = 193) samples, therefore requiring further validation.

**Figure 6:**
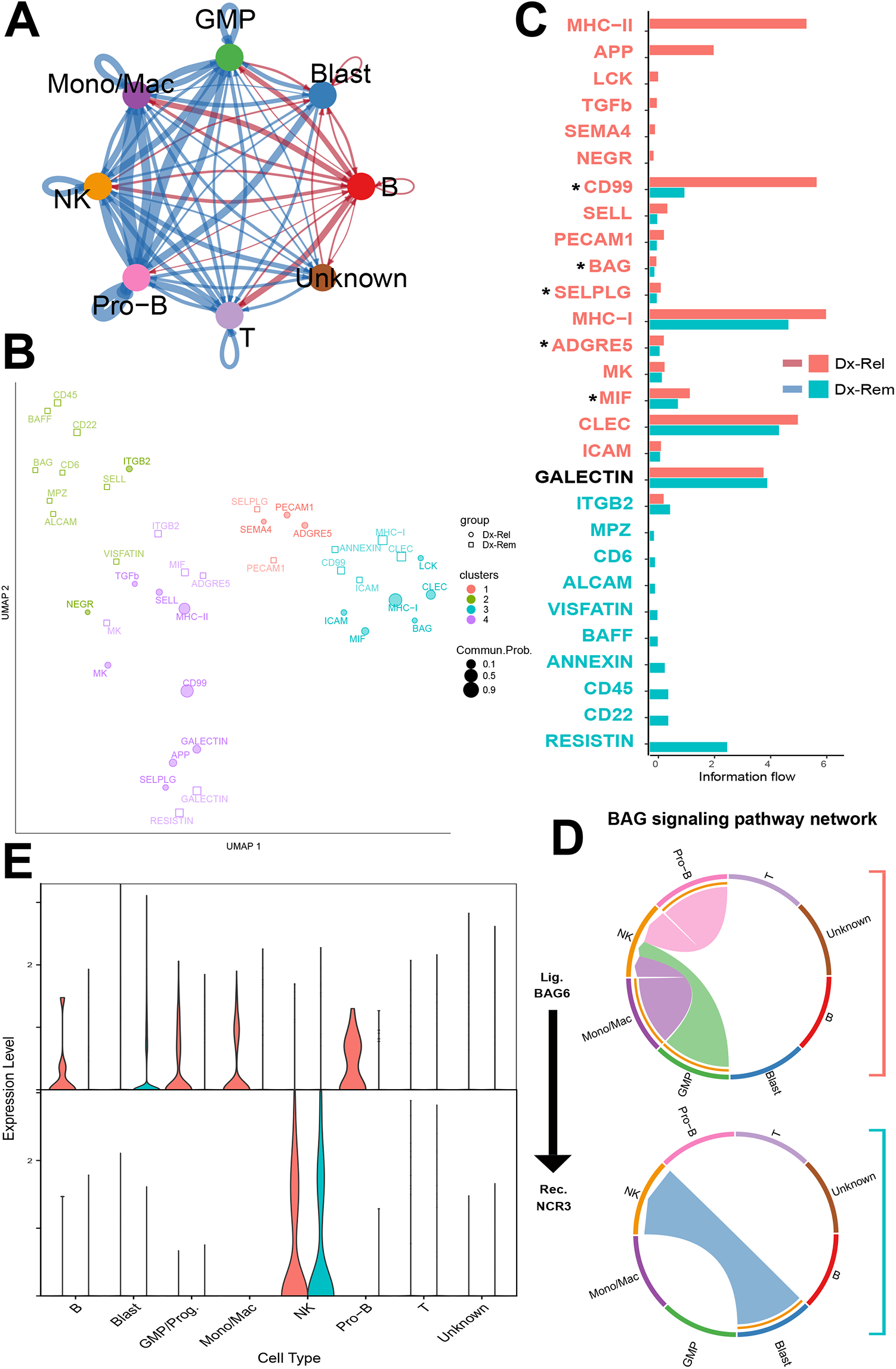
Cell communication analysis reveals enriched signaling pathways and ligand-receptor interaction differences between future relapse and remission groups. **A)** A circle plot showing the overall communication between cell types in Dx-Rel and Dx-Rem groups. The lines in the plot depict the communication network among the cell types. Lines are colored by the outcome groups (Red=Dx-Rel, Blue=Dx-Rem), with their thickness corresponding to the relative intensity of cellular communication measured based on ligand and receptor correlation. Dx-Rel and Dx-Rem groups show very different communication patterns between cell types. **B)** Clustering analysis of signaling pathways that were significantly enriched in each outcome group using manifold and classification learning analysis. The pathways form 4 major clusters with representation from Dx-Rel and Dx-Rem groups. Each cluster is represented with different colors. **C)** Comparison of signaling structure for individual ligands in Dx-Rel and Dx-Rem groups. Ligands in Dx-Rel and Dx-Rem samples were embedded based on the similarity of the sender and receiver cell types involved for said ligands. Ligands with similar sender and receiver cell types will have similar embeddings. A bar plot displaying the distance between Dx-Rel and Dx-Rem embeddings for each ligand is displayed. APP and CD99 have large differences in their pathway embedding. **D)** Detailed analysis of the BAG signaling pathway. Circos plots indicate the sender and receiver cell types involved in Dx-Rel and DX-Rem. Chords are colored by the sender cell type. BAG shows a very different signaling structure, with Blast cells as senders and NK cells as receivers in DX-rem only. In the case of Dx-Rel, Blast cells lack communication with most of the immune cells. **E)** Violin plot comparing the expression of the ligands (BAG6) and receptors (NCR3) between Dx-Rel (red) and Dx-Rem (green) samples across all cell types.

In addition, specific signaling pathways were estimated and functional similarities between the pathways inferred. The signaling pathways that were significantly enriched in each subset were compared using manifold and classification learning analysis and clustered based on their similarities (**Fig. 6B**). The dimensionality plot represents the overall functional similarity of the individual pathways for each clinical group. Cluster 1 consists of 5 pathways that include PECAM1, SEMA4, and ADGRE5 for the Dx-Rel group and SELPG and PECAM1 for the Dx-Rem group. These five signaling pathways have similar cell types and senders and receivers based on ligand and receptor expression, therefore, playing similar roles in the MPAL. In contrast, the CD99 pathway is significantly dysregulated in cluster 3 for the Dx-Rem group and cluster 4 for the Dx-Rel group indicating different functionality and association with outcomes (i.e., Dx-Rel and Dx-Rem). Further comparative analysis of the information flow or interaction strength of each signaling pathway in the Dx-Rem (blue bars) and Dx-Rel (red bars) groups is summarized in **Fig. 6C**. The CD99, BAG, SELPG, ADGRE5, and MIF signaling pathways were the top differentially enriched pathways (marked with an asterisk) based on their functional similarities/differences. To further explore differences in cellular contributions for the top 5 differentially expressed pathways, we generated chord diagrams showing the information flow and interactions among cell types for these pathways (**Fig. S8**). For example, the chord diagram for BAG signaling (**Fig. 6D**), shows that GMP, Mono/Mac, and Pro-B cells are the sender cell types for Dx-Rel samples, and blast cells are the sender cell type for Dx-Rem samples. In both cases, NK cells are the receiver cell type. The top ligand-receptor pair for this pathway found to be expressed in the samples is BAG6 to NCR3 (**Fig. 6E**). These results reveal that microenvironment cells have different communication machinery in the DX-Rel and DX-Rem groups even at baseline (disease diagnosis) that might be playing a role in disease progression and post-therapy clinical outcomes.

## Discussion

The emergence and optimization of single-cell profiling as a powerful tool to characterize the tumor microenvironment has revealed the heterogeneity of cancers, particularly different leukemia subtypes. Most MPAL biology studies to date, however, have focused on genetics and bulk RNA-sequencing profiling that measures an average signal from the amalgam of blast and immune microenvironment cells in the bone marrow, failing to address blast cell heterogeneity, blast and immune cell interactions, and the role of the immune microenvironment in clinical outcome. There has only been one study published utilizing a single-cell approach to analyze this rare leukemia. In this study, Granja *et al*. analyzed samples from five adult MPAL patients and compared their findings to controls for normal hematopoiesis (10). They demonstrated that despite widespread epigenetic heterogeneity within the patient cohort, common malignant signatures across patients were observed. Pediatric MPAL research is critical because in other leukemias like AML, significant differences have been demonstrated between adult and pediatric leukemia microenvironments (49-51). Also, the analysis by Granja *et al*. only included one patient with B/My MPAL; hence they did not perform a comparative analysis between the different B/My MPAL subtypes (10). Therefore, besides being the first study to characterize the single-cell tumor landscape in pediatric MPAL patients, our study is also the first study to compare single-cell expression profiles between the two major MPAL subtypes.

Comparative analysis of gene expression patterns showed that the two MPAL subtypes, B/My and T/My MPAL, had distinct single-cell transcriptomic profiles. The B/My MPAL cases had greater inter-patient heterogeneity and showed significant overlap with B-ALL and to a lesser extent with AML. The T/My MPAL cases, on the other hand, had less inter-patient variability but formed a single distinct expression profile unique from the other leukemia subtypes. These results support the findings from the large pediatric MPAL genomic study published in 2018, in which Alexander *et al*. showed that B/My MPAL and T/My MPAL had distinct genetic profiles based on transcriptome and whole-genome sequencing (7). B/My MPAL shared common genomic features with both B-ALL and AML, and T/My MPAL showed a similar mutational profile to early T-cell precursor ALL (ETP-ALL) (52). These findings suggest that the two MPAL subtypes should be considered distinct entities; this may have implications for differing treatment regimens if validated in future studies. While there is no clear consensus as to how to treat MPAL patients, more recent literature has suggested utilizing an ALL-directed therapy approach. While our results do support this treatment approach for B/My MPAL given the overlap with the B-ALL gene expression profile, the T/My MPAL treatment regimen may need to be re-considered based on its unique profile compared to other leukemias.

Genes that were overexpressed in the blasts of both MPAL subtypes compared to other acute leukemias included *CD81* and *UBE2S*. CD81 has been associated with a poor prognosis in AML (53) and is also a known marker in B-ALL (54, 55) while the E2 family of a ubiquitin-conjugating enzyme, UBE2s has a role in the cell cycle progression (56). AML cells are dependent on UBE2N-dependent oncogenic immune signaling states (57). DEG-associated pathway analysis showed that Sirtuin signaling, p38 MPAK signaling, and PI3K signaling were upregulated in B/My MPAL blasts. Sirtuin signaling genes such as *SIRT1, SIRT2*, and *SIRT6* promote B-ALL, AML, and T-ALL leukemia cell survival, and *SIRT1* is proposed as an unfavorable prognosis marker in the adult AML (32, 58-60). In T/My MPAL blasts, oxidative phosphorylation, and Rho family GTPases signaling were upregulated, known to promote leukemia cell survival (35).

Single-cell profiling of the non-blast bone marrow microenvironment cells showed MPAL B cells overexpressed *IGHG1, JCHAIN*, and *IGHG3* and cytotoxic T cells overexpressed *NFKBIA, SYNE2*, and *COTL1*. Pathway analysis of the microenvironment cells showed that in B/My MPAL, T cells were involved in CRH signaling, NGF signaling, and the AP1 pathway. Studies show that NGF regulates T cell proliferation (61), and that exhausted T cells have low expression of AP-1, which is a regulator of T cell activation (62). The upregulation of these pathways in B/My MPAL T cells suggests that there may be activity related to immunosuppression, T cell proliferation, and T cell activation present. Enrichment of the FOSB pathways in B/My MPAL B cells and the TNFA signaling pathway via NF-κB in progenitor cells have been associated with poor leukemia outcomes (45). For T/My MPAL, B-lymphoid cells showed enrichment of multiple pathways including IL-10 signaling, the CSK pathway, and IFN-γ signaling. IL-10 is an immune-suppressive cytokine and while expressed in B cells has been associated with the inhibition of pro-inflammatory cytokines (63). Because IL-10 signaling is uniquely upregulated in T/My MPAL B cells, this suggests there is a more immune-suppressive environment than in other leukemias.

Finally, more recent literature has shown that pediatric MPAL patients with MRD at EOI have significantly poorer outcomes (4, 64). In the multi-national iBFM-AMBI2012 study, Hrusak *et al*. showed that patients with MRD at EOI had a significantly worse event-free survival (EFS) and overall survival (OS), despite their analysis being complicated by the inclusion of a myriad of treatment regimens (4). A more recent multicenter study by Oberley *et al*. on MPAL showed that MRD positivity was highly predictive of relapse and death, with significantly worse EFS and OS among MRD-positive patients. Based on these findings, we performed a comparative analysis of our diagnostic samples based on EOI MRD and future remission or relapse status. Although we had a limited number of samples, our data suggest that unique transcriptome profiles at diagnosis can be associated with MRD at EOI as well as future relapse. Thus, identifying specific gene expression profiles associated with MRD positivity and relapse would allow for better risk stratification and more tailored therapy for MPAL in future prospective clinical trials.

## Conclusions

Our data provide an initial framework of the single-cell landscape of pediatric MPAL. Gene signatures and pathways specifically enriched in B/My and T/My MPAL subtypes were identified. While a larger sample size is needed to validate these findings, these signatures could be used to identify potential targets for the development of novel diagnostics and therapies in the future.

## Supplementary Figure Legends

**Figure S1: Workflow for MPAL biomarker identification steps**. The flowchart diagram shows the filtering steps performed to identify the Mixed Phenotype Acute Leukemia (MPAL) subtype biomarkers in **Fig. 2** and **Fig. 4**. Each oval in the flowchart represents a yes/no decision, if the gene does not meet the requirements (N: No) it is removed from the potential biomarker list and if the gene does meet the requirements (Y: Yes) it moves on to the next analysis step. Step 1: consists of the three filtering criterions (p-value < 0.05, log2FC > 0.25 or Fold change >1.2, and percent cells expressing gene > 50%). Step 2: filtering out genes with >0.5 average expression levels in healthy immune clusters of the Human Cell Atlas (HCA) (21). Step 3: filtering out genes with >0.5 average expression levels among HCA hematopoietic stem cells (HSCs). Step 4: checking gene expression by feature plots (little to no expression in feature plots of HCA immune cells and HSCs, see **Figure S3** for examples).

**Figure S2: Expression profile of canonical cell-type markers in MPAL and healthy BM samples**. The dot plot shows the expression of canonical cell-lineage markers in the different cell types from MPAL, and young adult healthy BM samples shown in **Fig. 1**. The dot size represents the percentage of cells expressing a specific gene, and the color represents its level of expression with red and blue colors showing high and low expression, respectively.

**Figure S3: HCA feature plots showing expression of MPAL biomarker genes**. The feature plots show the expression of the different MPAL biomarkers in the Human Cell Atlas (HCA) (21) immune cells (**Fig. S3A**) and hematopoietic stem cells (**Fig. S3B**). The cells are colored on a color scale from grey (low) to dark purple (high) based on the gene expression.

**Figure S4: Pathways enriched in B/My and T/My MPAL blast cells**. The bar plot shows the common significantly enriched pathways (P-value <.05) of B/My and T/My MPAL blasts on the Y-axis and the Z-score representing potential activation or inhibition on the X-axis. A single asterisk (*) next to a pathway name represents whether the pathway is associated with promoting leukemia in the literature, and two asterisks (**) represent a pathway that is associated with inhibiting leukemia. The z-scores of B/My MPAL blasts are shown in orange, and T/My MPAL is shown in purple. The analysis was performed using the Ingenuity pathways analysis platform (Qiagen Inc.)

**Figure S5: Comparative analysis of cellular communication among different leukemias**. The comparative analysis of cellular communication was performed among different leukemias and healthy bone marrow samples to identify pathways showing different communication in MPAL. The bar plots show the relative (left) and absolute (right) information flow. Information flow (x-axis) represents the sum of communication probability for each cell type for specific signaling pathways (y-axis). For example, T-ALL (blue) microenvironment cells have the highest communication probability in the Selectin P Ligand (SELPLG) signaling pathway. See **Figure S8** for information on the specific cell type communication for the MPAL top signaling pathways.

**Figure S6: Expression of canonical cell type markers in leukemia and healthy bone marrow samples**. The dot plot shows the expression of canonical cell-lineage markers in the different cell types present in the annotated leukemia and healthy sample microenvironment shown in **Figure 3**. The dot size represents the percentage of cells expressing a specific gene, and the color represents its level of expression with red and blue colors showing high and low expression respectively.

**Figure S7: Kaplan-Meier survival curves for B/My Dx-Rel and Dx-Rem genesets**. Survival analysis was performed on bulk RNA-seq data from B/My MPAL samples in the TARGET-ALL-P3 (ambiguous leukemia) dataset. The samples were split into high (red) and low (blue) expression groups based on the enrichment values calculated by performing gene set enrichment analysis using the gene sets from the Dx-Rel (future relapse) and Dx-Rem (future remission) groups. Survival of the high and low expression groups was calculated and plotted using the survival and survMisc packages in R. The Dx-Rel (**A**) gene set survival analysis resulted in a logRank p-value of 0.054 and Cox hazard ratio (HR) and p-value of 0.17 and 0.091. The Dx-Rem (**B**) gene set survival analysis resulted in a logRank p-value of 0.326 and Cox HR and p-value of 2.17 and 0.338, respectively.

**Figure S8: CellChat chord diagrams for top differentially enriched signaling pathways in MPAL Dx-Rel and Dx-Rem samples**. CellChat analysis was performed on the MPAL Dx-Rel (future relapse) and Dx-Rem (future remission) samples to determine the differences in cell type interactions between the two groups. Functional similarity was calculated to identify the top differentially enriched signaling pathways between the two groups (ADGRE5, BAG, MIF, SELPLG, and CD99). The resulting chord diagrams are shown for each signaling pathway (**A-E**). The arrows show the information flow from the predicted sender to the receiver cell types in Dx-Rel (future relapse) and Dx-Rem (future remission) groups.

## Supplementary Table Legends

**Table S1. List of top 10 upregulated genes in MPAL microenvironment cell types as compared to corresponding cells in healthy bone marrow samples**.

**Table S2. List of significantly differentially expressed genes in MPAL as compared to healthy control and blast cells from other leukemias. A)** The table lists genes that are significantly differently expressed in either MPAL or specific subtype (i.e., B/My MPAL, T/My MPAL) blast cells as compared to healthy control and blast cells from other leukemias based on Wilcoxon rank test estimated P-Val < 0.05, log2FC > 0.25, and percent expression in the target group(pct.1) > 0.5. **B)** List of B/My MPAL, T/My MPAL, and MPAL blast markers after filtering out genes with >0.5 average expressions in clusters of the HCA (21) healthy immune cell dataset. **C)** List of B/My MPAL, T/My MPAL, and MPAL blast markers after filtering out genes with >0.5 average expressions in stem cell clusters of the HCA healthy hematopoietic stem cell (HSC) dataset. **D)** List of B/My MPAL, T/My MPAL, and MPAL blast markers that show minimal expression in any immune cell and stem cell clusters of the HCA datasets. **E)** List of genes that show no patient-specific heterogeneity in MPAL samples determined using visual analysis of heatmaps.

**Table S3. The top upregulated signaling pathways for MPAL blasts**. The significantly enriched (P-value <.01) and activated (Z score ≥1) signaling pathways for **A)** B/My and **B)** T/My MPAL blasts. The table contains the pathway names, the -log(p-value), the z-score, and the genes.

**Table S4. Gene sets associated with MPAL remission or relapse outcomes**. The gene sets were identified based on differential expression analysis between remission and relapse samples for each MPAL subtype. The significance of the differentially expressed genes was determined based on the Wilcoxon rank test based on P-value and fold change (P-value < 0.01 and average log2FC > 1.25).

