## Supplementary figures and images for "Single Cell RNA Sequencing Driven Characterization of Pediatric Mixed Phenotype Acute Leukemia"

### Supplemental Figures

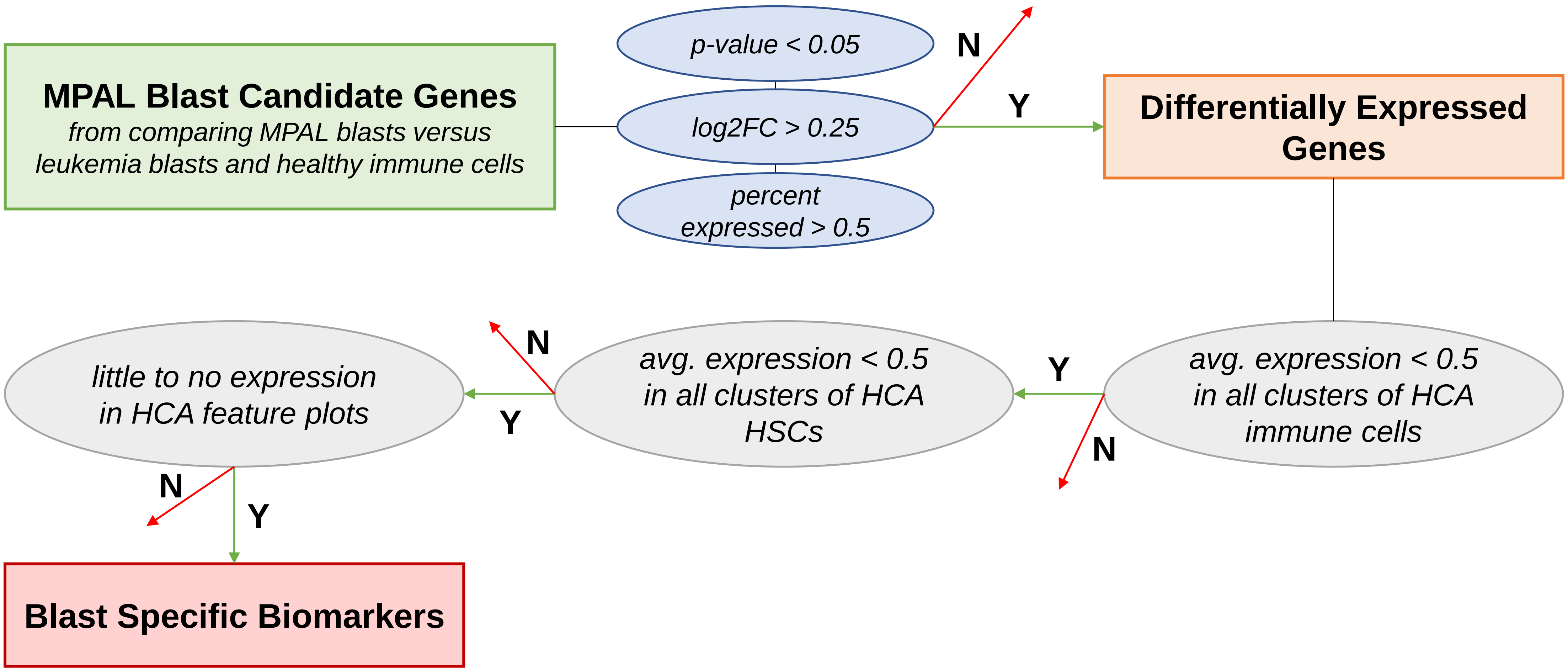

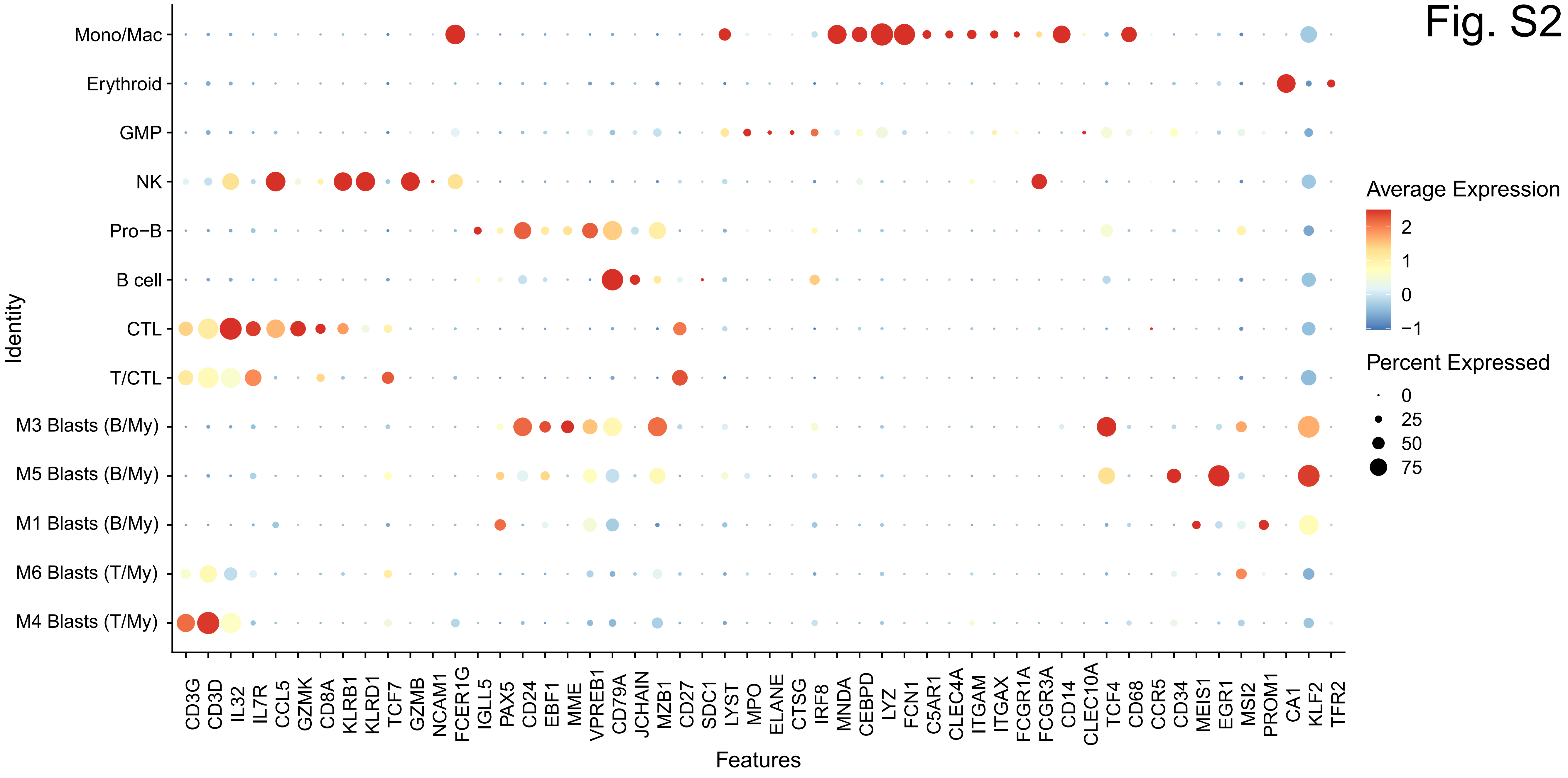

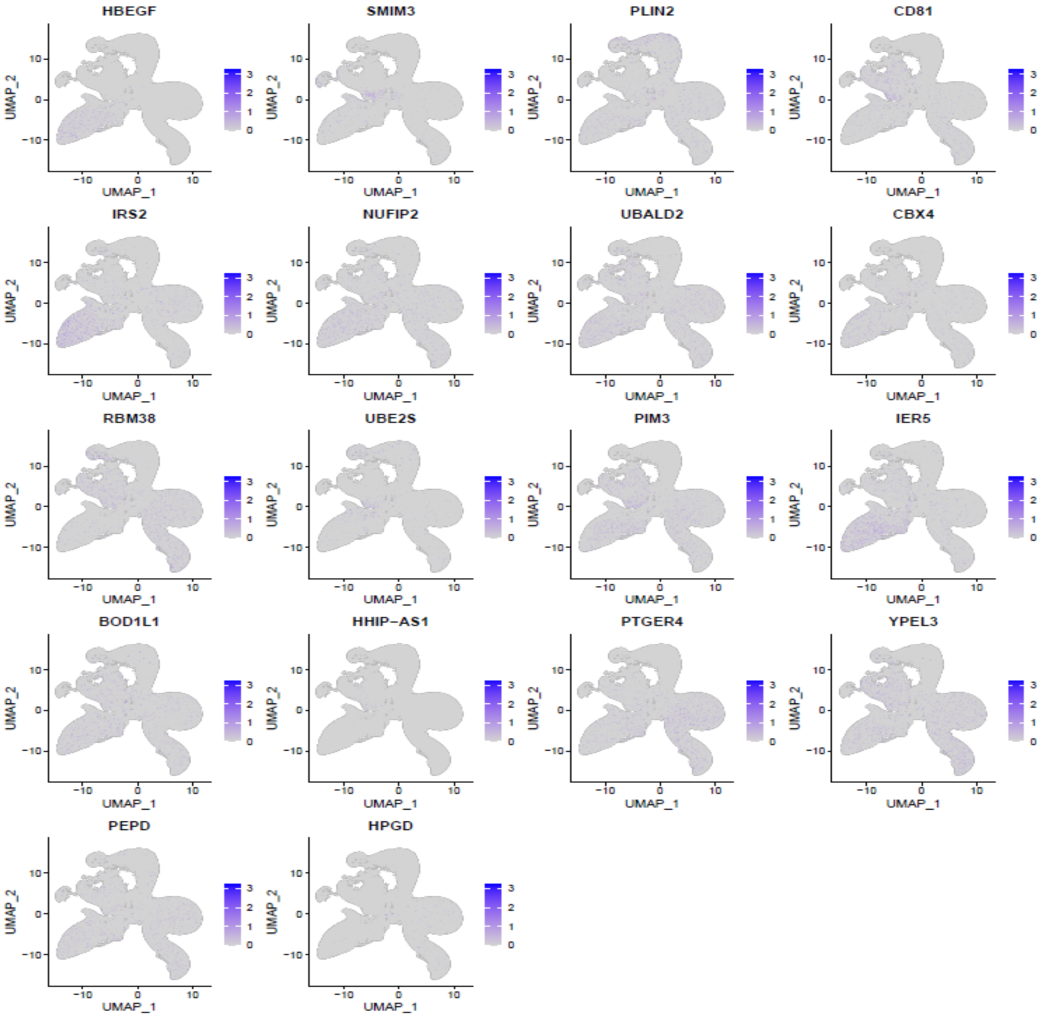

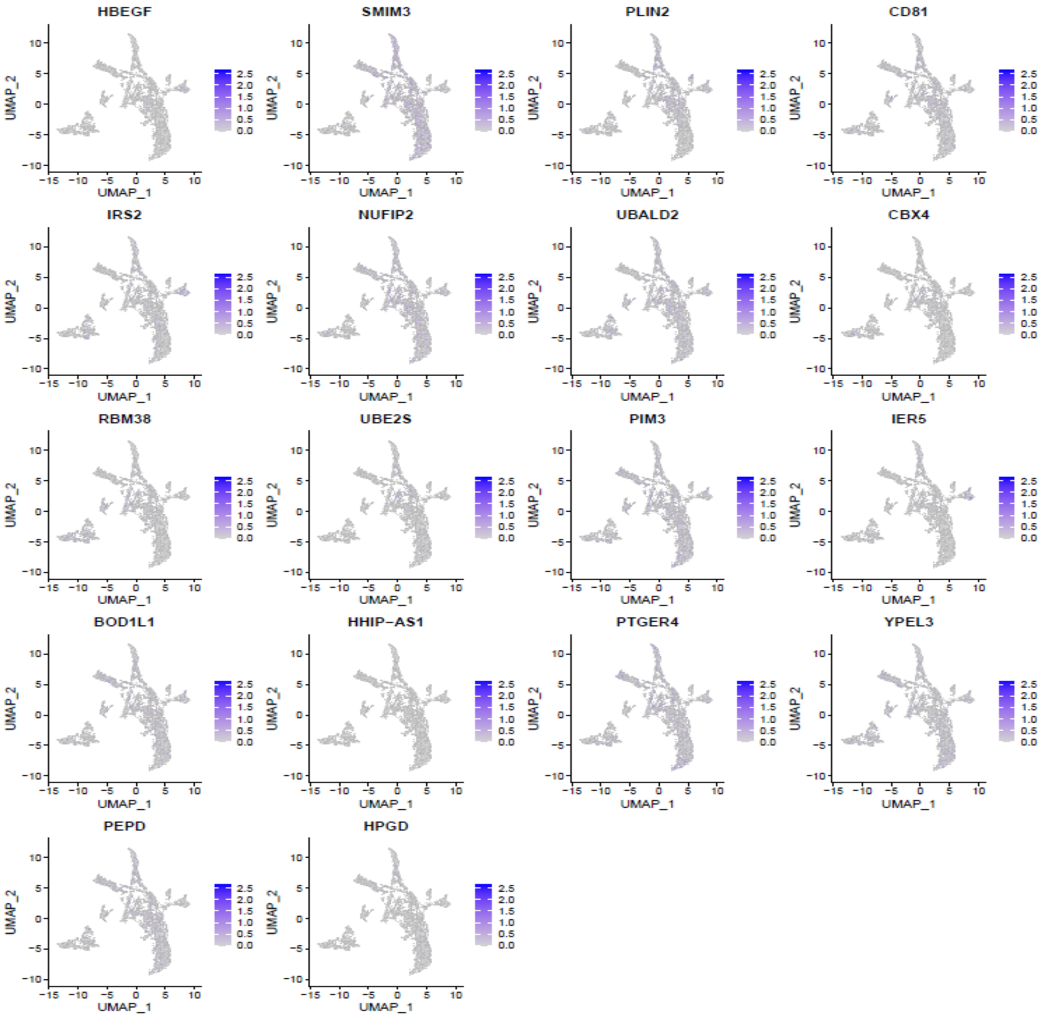

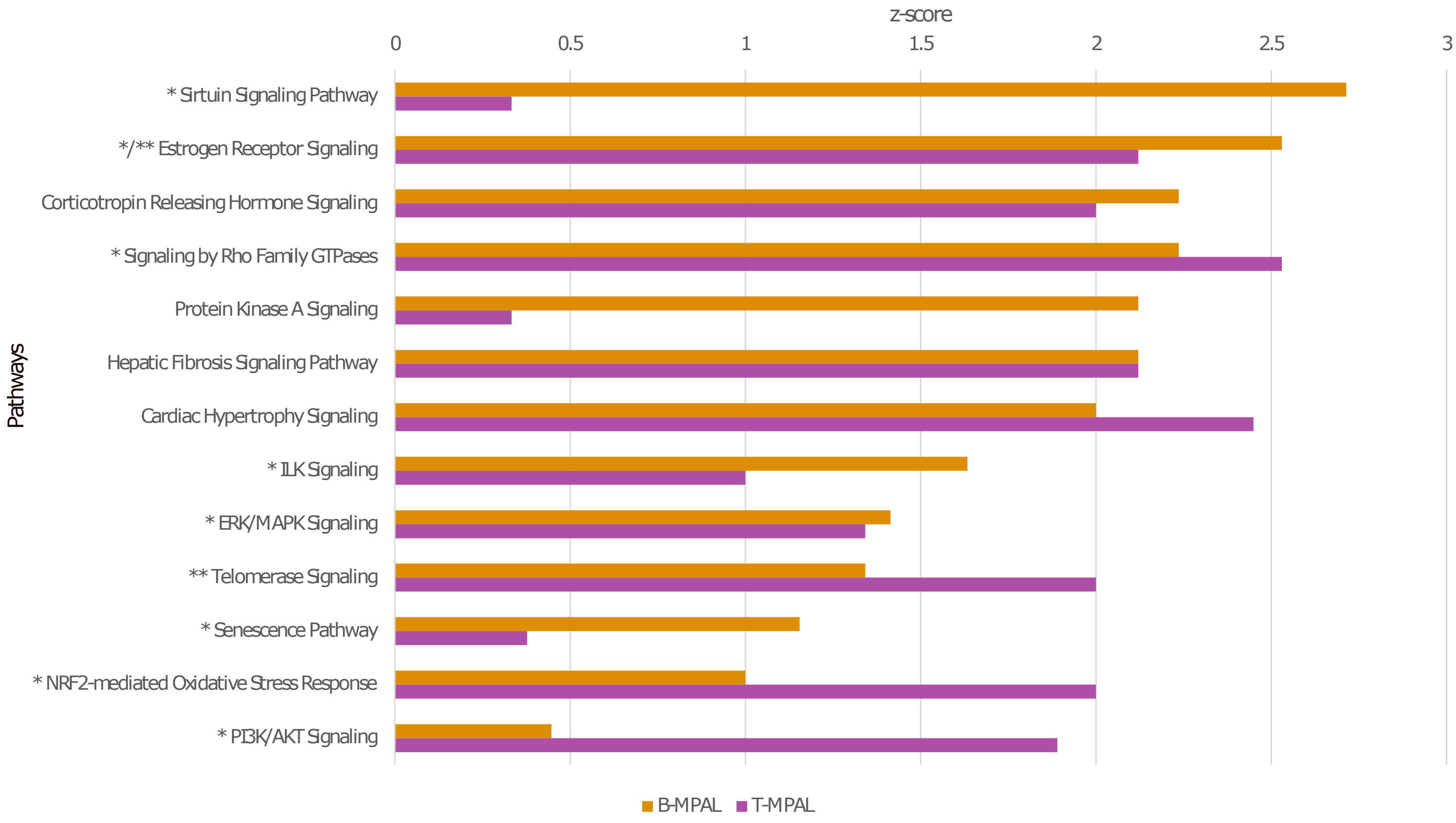

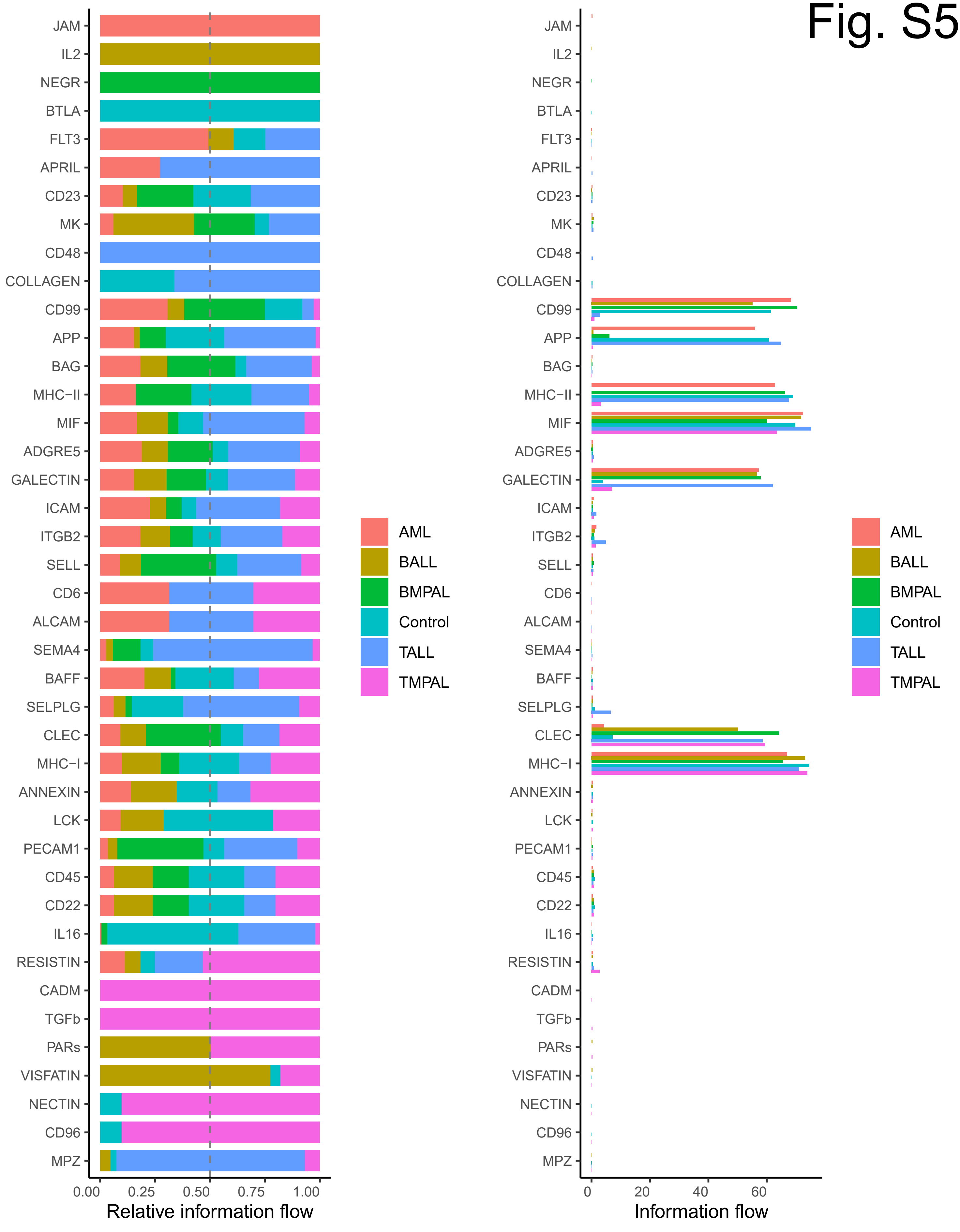

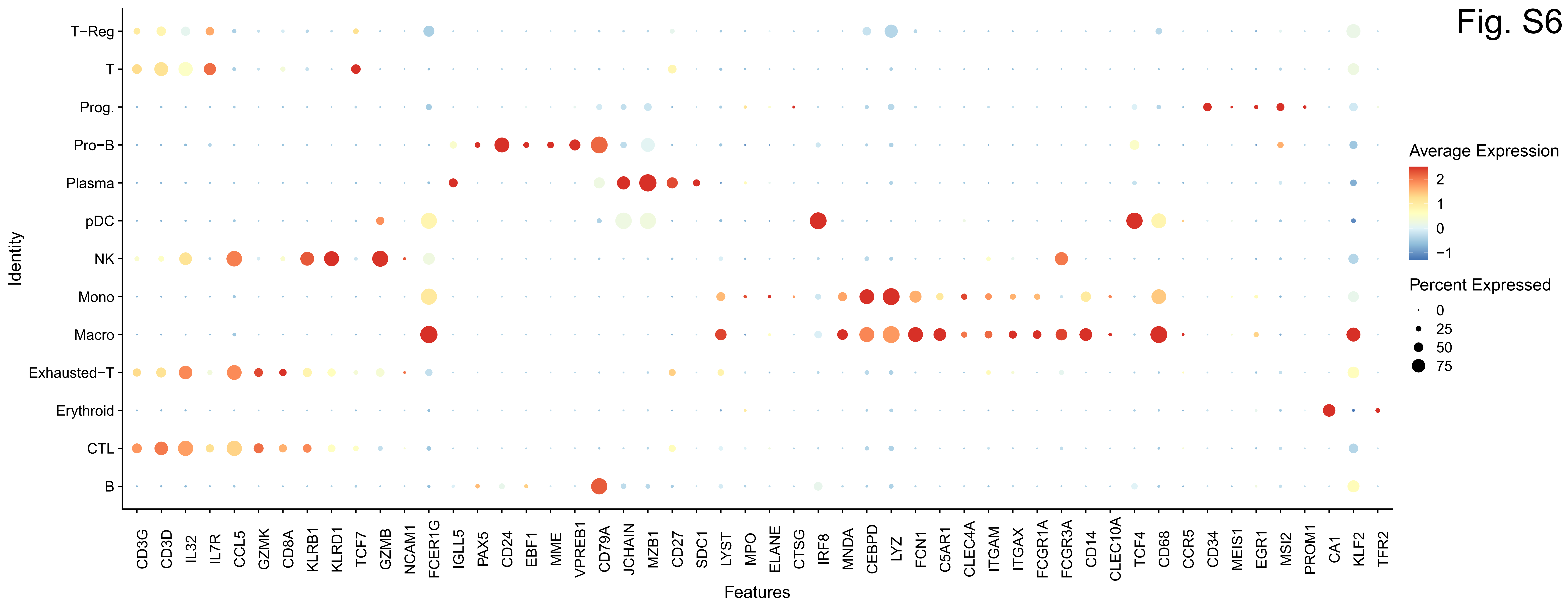

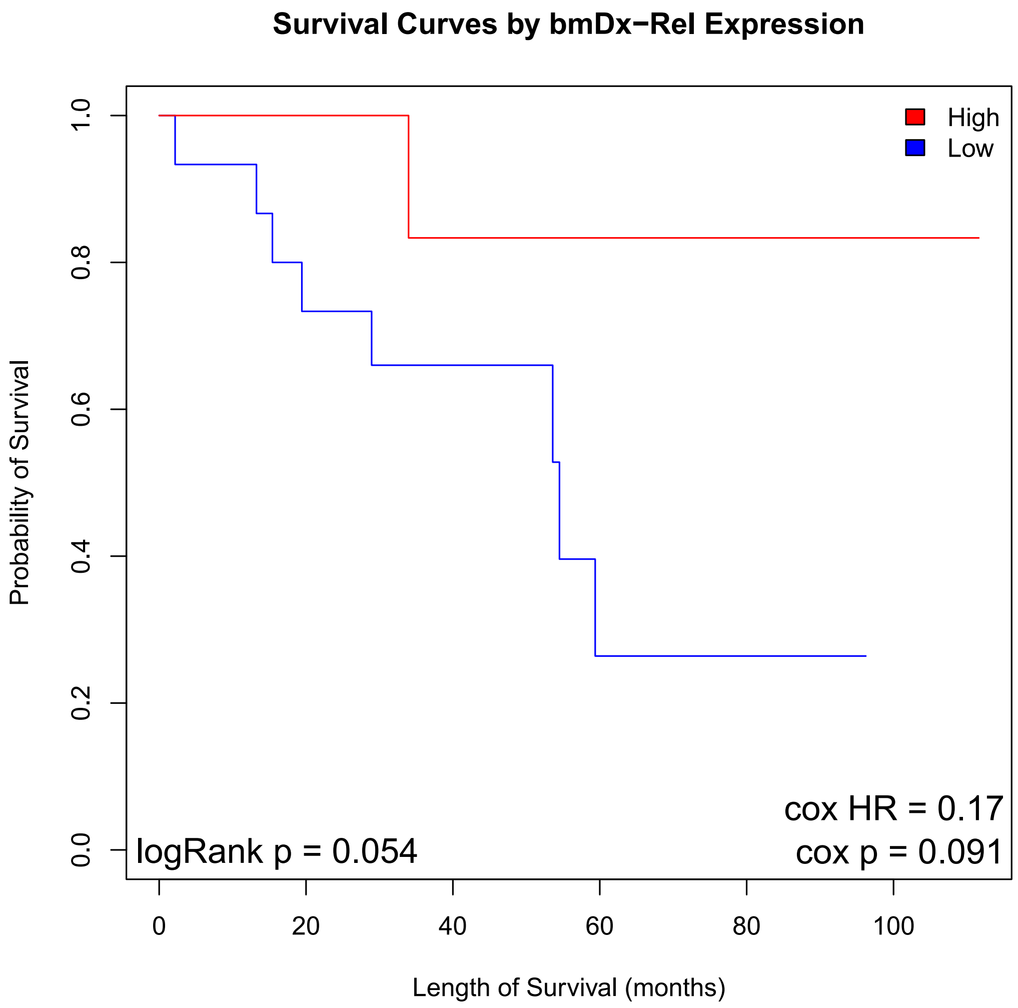

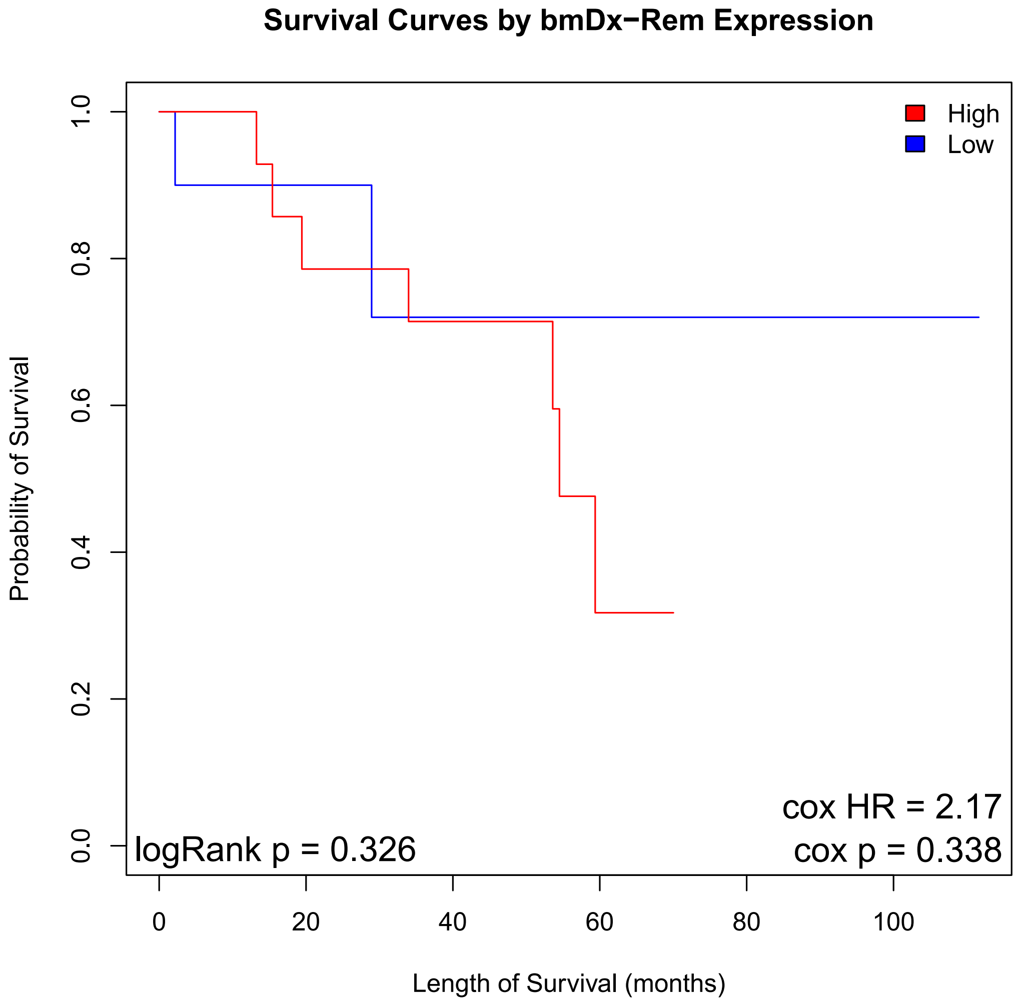

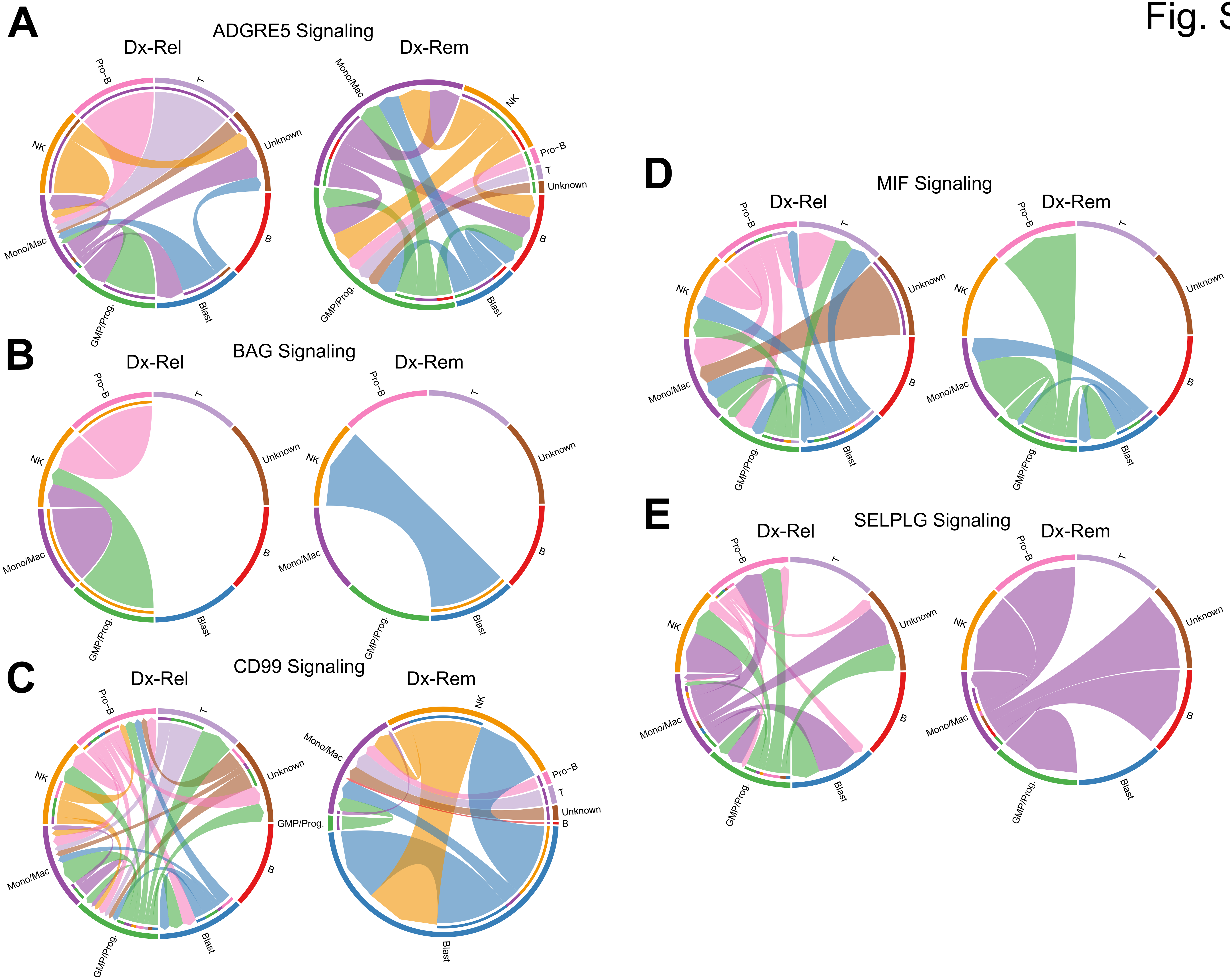
